# Broad and potent neutralizing mAbs are elicited in vaccinated individuals following Delta/BA.1 breakthrough infection

**DOI:** 10.1101/2023.03.30.534872

**Authors:** Jeffrey Seow, Zayed A. Shalim, Carl Graham, Simon Kimuda, Aswin Pillai, Thomas Lechmere, Ashwini Kurshan, Atika M. Khimji, Luke B. Snell, Gaia Nebbia, Christine Mant, Anele Waters, Julie Fox, Michael H. Malim, Katie J. Doores

**Author notes:** These authors contributed equally (Jeffrey Seow, Zayed A. Shalim).

## Abstract

Despite the success of COVID-19 vaccines in preventing infection and/or severe disease, with the emergence of SARS-CoV-2 variants of concern (VOC) which encode mutations in Spike, and the waning of vaccine induced immunity, there has been an increase in SARS-CoV-2 infections in vaccinated individuals which leads to increased serum neutralization breadth. However, how exposure to a heterologous Spike broadens the neutralizing response at the monoclonal antibody (mAb) level is not fully understood. Through isolation of 119 mAbs from three individuals receiving two-doses of BNT162b2 vaccine before becoming delta or omicron/BA.1-infected, we show that breadth arises from re-activation and maturation of B cells generated through previous COVID-19 vaccination rather than a *de novo* response specific to the VOC Spike. Isolated mAbs frequently show reduced neutralization of current circulating variants including BA.2.75.2, XBB, XBB.1.5 and BQ.1.1 confirming continuous selective pressure on Spike to evolve and evade neutralization. However, isolation of mAbs that display effective cross-neutralization against all variants indicate the presence of conserved epitopes on RBD and a lesser extent NTD. These findings have implications for selection of Spike antigens for next-generation COVID-19 vaccines.

## Introduction

Both SARS-CoV-2 infection and COVID-19 vaccines based on the SARS-CoV-2 surface glycoprotein, Spike, generate neutralizing antibodies in SARS-CoV-2 naïve individuals which can prevent infection and/or severe disease. Indeed, induction of neutralizing antibodies is a correlate of protection^1–4^. Through isolation of monoclonal antibodies (mAbs) from SARS-CoV-2 convalescent donors or COVID-19 vaccinees, we and others have identified several neutralizing epitopes on Spike^5–12^, including epitopes on the receptor binding domain (RBD), N-terminal domain (NTD), S1D domain of S1, and on S2. mAbs against many of these epitopes have been shown to protect from SARS-CoV-2 infection in animal challenge models^13–15^.

However, with the waning of vaccine induced immunity^16, 17^ and the emergence of SARS-CoV-2 variants of concern (VOC) which encode mutations in Spike^18^, there has been an increase in infections with VOCs in vaccinated individuals. We and others have previously shown that a breakthrough infection (BTI) with a VOC following vaccination can broaden the neutralization capacity of the polyclonal response in sera, and generate neutralizing activity against highly divergent SARS-CoV-2 viral variants carrying Spike mutations across multiple neutralizing epitopes^19–23^. Despite the increase in infections with new VOCs, vaccines based on the ancestral SARS-CoV-2 (Wuhan-1) have remained effective at reducing severe disease and hospitalizations^24, 25^. For continued control of the SARS-CoV-2 pandemic, it is important to understand how infection with SARS-CoV-2 variants in vaccinated individuals shapes the antibody response against SARS-CoV-2 Spike and the resulting susceptibility to infection with newly arising VOCs. Further understanding in this area has direct application to selecting Spike antigens to be used in future generation COVID-19 vaccines.

In the context of influenza, secondary infection with an antigenically distinct influenza strain generates antibodies that are highly cross-reactive with the primary infecting virus (termed original antigenic sin or immune imprinting)^26–28^. This is thought to arise due to preferential induction of antibodies with higher affinity to the priming antigen than the boosting antigen. A third COVID-19 vaccine dose based on the Wuhan-1 Spike has also been shown to increase neutralization breadth against VOCs, in particular against omicron/BA.1^8, 19, 29, 30^. However, whether a SARS-CoV-2 variant infection in vaccinated individuals leads to a *de novo* response specific for the infecting VOC or whether pre-existing memory B cells are activated upon VOC exposure is not fully understood.

Here, we isolated mAbs from three individuals who had received two doses of the BNT162b2 vaccine and then experienced a delta or omicron/BA.1 infection to understand how neutralization breadth increases following BTI at the mAb level. We used antigen-specific B cell sorting with an S1 probe matching the vaccine and infecting variant to isolate 119 mAbs. We show that all isolated mAbs can bind and neutralize vaccine and infection strains, with the majority of neutralizing mAbs targeting the RBD indicating re-activation and continued maturation of B cell clones generated through previous COVID-19 vaccination. Isolated mAbs showed strong cross-neutralization of omicron sub-lineages BA.1, BA.2 and BA.4/5 but the majority showed reduced neutralization against newer variants, including BA.2.75.2, XBB, XBB.1.5 and BQ.1.1. However, subsets of mAbs with broad cross-neutralization were identified highlighting the presence of conserved neutralizing epitopes across antigenically distant Spikes. These findings have implications for selecting Spike antigens for next generation COVID-19 vaccines.

## Results

### Wuhan-1 and VOC S1 reactive B cells present at similar levels

To gain insight into the neutralizing activity within polyclonal sera from BTI individuals, we used antigen-specific B cell sorting to isolate S1-reactive IgG+ B cells from two Delta-infected individuals (VAIN1 and VAIN2) and one BA.1-infected individual (VAIN3) (see **Supplementary Figure 1** for full sorting strategy). All three donors had no history of SARS-CoV-2 infection and had received 2-doses of the BNT162b2 vaccine with an extended interval^19^ prior to infection. Blood samples were collected 14, 87 and 26 days post infection, respectively (See **Supplementary Table 1** for full donor information). Cross-neutralizing activity was observed in sera collected at these time points (**Supplementary Figure 2**). To allow for identification of variant specific mAb responses, we performed two sorts from each donor using different antigen-baits, one sort using the Wuhan-1 S1 (matched vaccine strain and referred to as wild-type, WT) and one sort using the VOC S1 (delta S1 for VAIN1 and VAIN2, and BA.1 S1 for VAIN3) (**Figure 1A**). Similar levels of WT and VOC reactive IgG^+^ B cells were observed for all three donors (**Figure 1B**).

**Figure 1.**
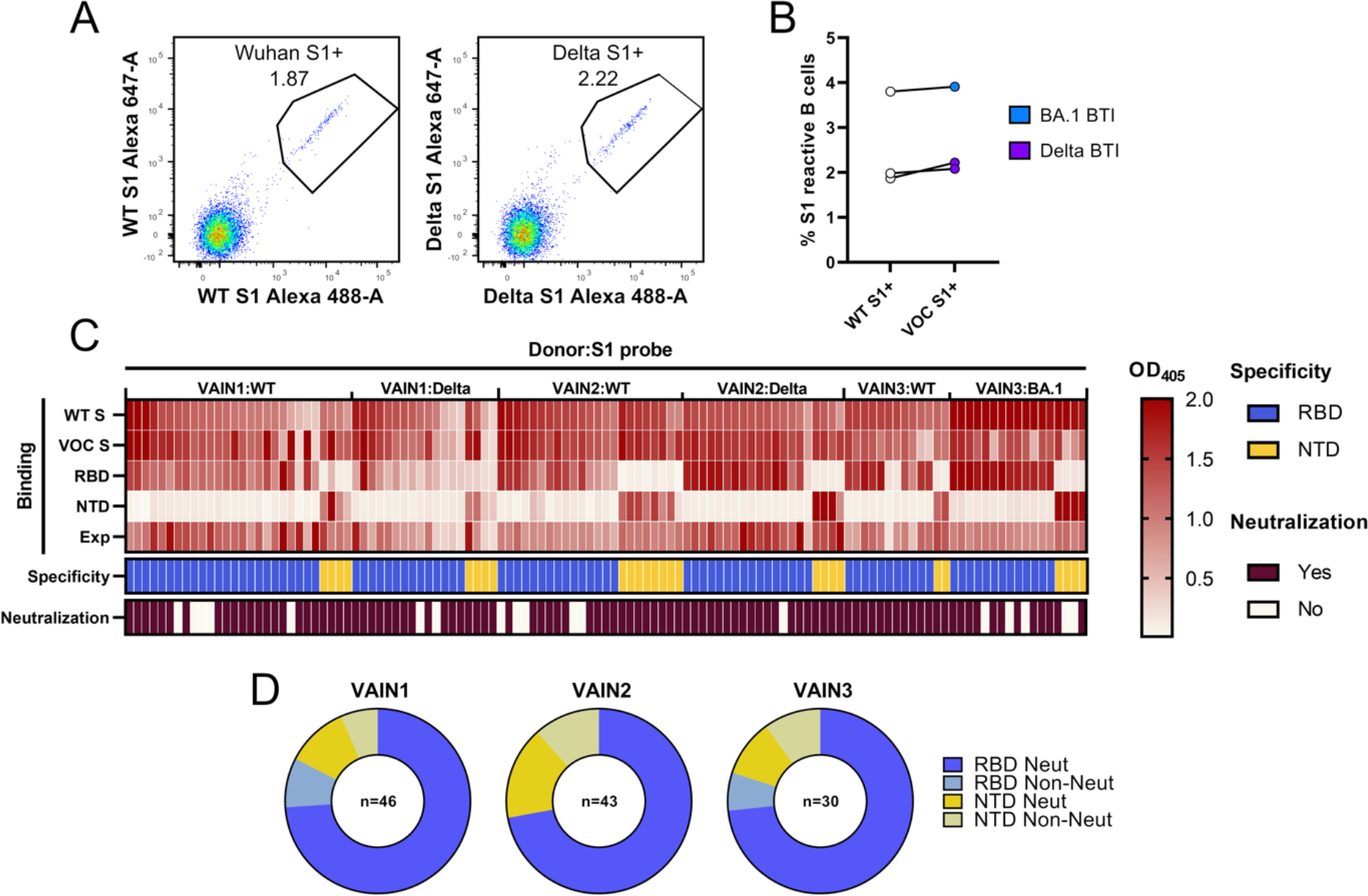
Isolation of mAbs using antigen-specific B cell sorting. A) CD14^-^/CD3^-^/CD8^-^/CD19^+^/IgM^-^/IgD^-^/IgG^+^ and S1^+^ B cells were sorted into 96-well plates. Example fluorescent activated cell sorting (FACS) showing percentage of CD19^+^IgG^+^ B cells binding to S1 of Wuhan-1 or S1 of delta VOC. Full sorting gating strategy is shown in **Supplementary** **Figure 1**. **B)** Percentage of CD19^+^IgG^+^ S1 Wuhan and S1 VOC reactive B cells for each donor (delta for VAIN1 and VAIN2, BA.1 for VAIN3). Data points from the same individuals are linked. Blue: BA.1/Omicron infected, purple: delta-infected. **C)** Heatmap showing IgG expression level and binding to SARS-CoV-2 Spike (WT and VOC (delta for VAIN1 and VAIN2, BA.1 for VAIN3), and to Spike domains RBD and NTD. The figure reports OD values from a single experiment (range 0–2.0) for undiluted supernatant from small-scale transfection of 119 cloned mAbs. Antigen binding was considered positive when OD at 405 nm was >0.2 after subtraction of the background. SARS-CoV-2 Spike domain specificity (RBD or NTD) for each antibody is indicated. Neutralization activity was measured against wild-type (WT; Wuhan) pseudotyped virus using concentrated supernatant and neutralization status is indicated. Antigen probe used to select specific B cells is indicated (i.e. WT S1, delta S1 or BA.1 S1). **D)** Distribution of mAbs targeting RBD and NTD for each donor, as well as their neutralization capability. mAbs are classified as shown in the key.

mAb heavy and light chain genes were rescued using reverse transcription and nested PCR using gene-specific primers^31, 32^. The variable regions were then cloned into IgG1 expression vectors using Gibson assembly and directly transfected in the HEK293T/17 cells^5, 6^. Crude supernatants were tested by ELISA and the heavy and light chain genes of Spike reactive IgGs were sequenced. In total, 46, 43 and 30 spike-reactive mAbs were isolated from VAIN1, VAIN2 and VAIN3, respectively (**Figure 1C**).

### Delta and BA.1 BTI generates neutralizing mAbs against RBD and NTD

ELISA with the crude supernatants were used to determine the VOC specificity and the specific domains targeted by each mAb. Despite different antigen-baits being used for B cell selection, all mAbs isolated showed reactivity to both the WT and VOC Spikes, consistent with reactivation of B cells generated from prior vaccination (**Figure 1C**). Similar to previous observations^5, 6^, 72.1-83.3% of mAbs were RBD specific (**Figure 1D**) with the remaining mAbs specific for NTD.

Neutralization activity of concentrated supernatant was determined using HIV-1 virus particles, pseudotyped with SARS-CoV-2 Wuhan-1 (wild-type, WT) Spike^33^. As previously observed, the majority (93.5 %) of RBD-specific mAbs had neutralizing activity (**Figure 1D**) whereas only 53.8 % of NTD mAbs showed neutralizing activity against WT pseudotyped virus.

### Mutation and germline gene usage

The level of somatic hypermutation and germline gene usage was determined using the IMGT database^34^. The mean divergence from germline at the nucleotide level for the variable heavy (V_H_) and light (V_L_) regions was 5.0% and 3.9%, respectively (**Figure 2A**). Comparison of mutation levels between the three donors showed that VAIN3 (BA.1-infected) had higher mutation than VAIN1 and VAIN2 in the V_H_ and V_L_ region (**Supplementary Figure 3A**). mAbs selected using the BA.1 S1 probe were more mutated than delta or WT S1 selected B cells (**Supplementary Figure 3B**). However, this might be donor specific observation as there was no difference in the level of mutation in V_H_ between WT S1 and VOC S1 selected B cells from each donor (**Supplementary Figure 3C**).

**Figure 2:**
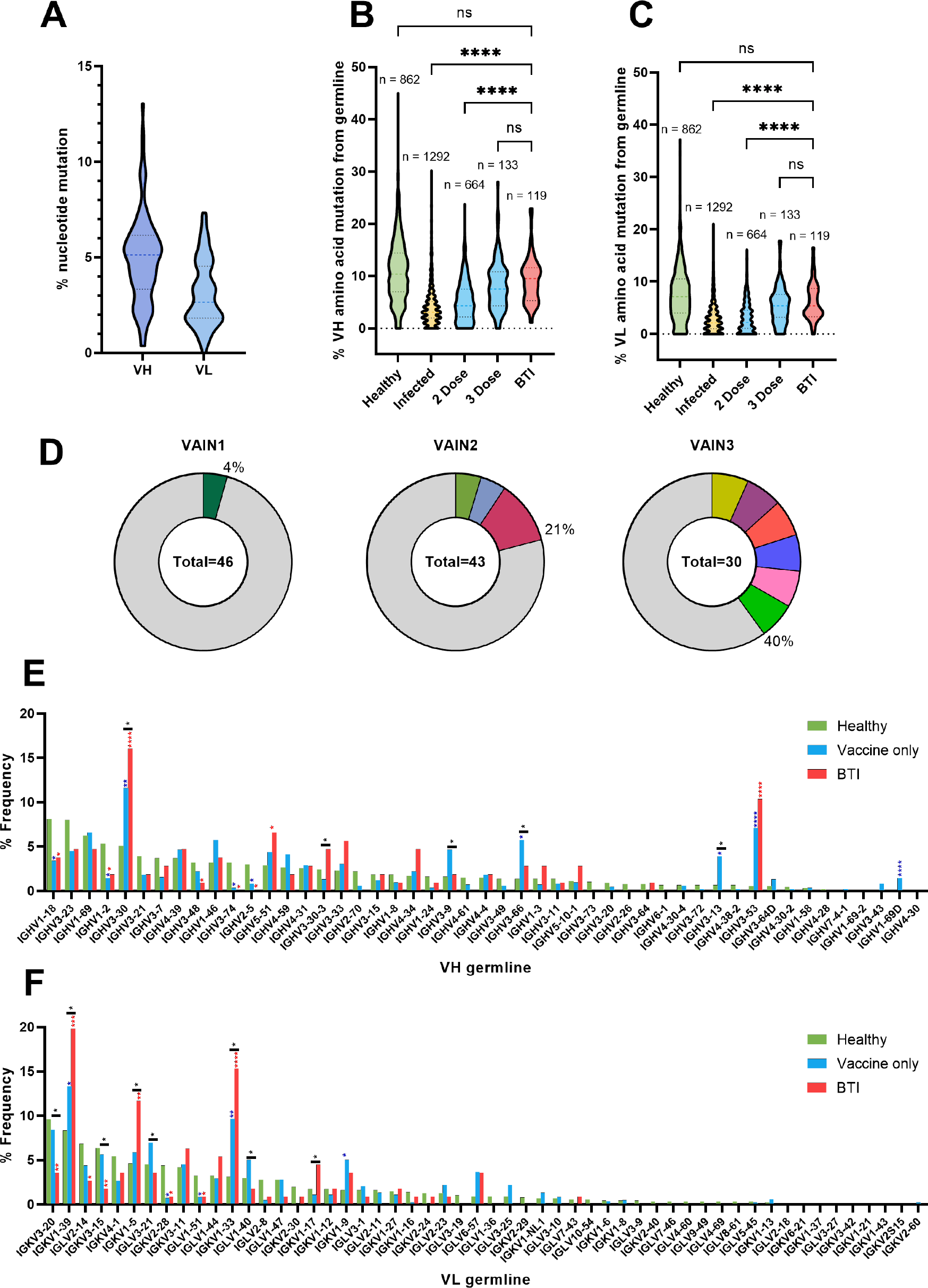
BTI mAbs show higher somatic hypermutation than mAbs isolated following two vaccine doses. A) Truncated violin plot showing the percentage of nucleotide mutation compared with germline for the VH and VL genes of Spike-reactive mAbs isolated from VAIN1, VAIN2 and VAIN3. Truncated violin plot comparing the percentage of amino acid mutation compared with germline for **B)** VH and **C)** VL between Spike-reactive mAbs isolated following infection, 2 doses of vaccine, 3 doses of vaccine or following BTI and IgG BCRs from SARS- CoV-2-naive individuals^36^. D’Agostino and Pearson tests were performed to determine normality. Based on the result, a Kruskal-Wallis test with Dunn’s multiple comparison post hoc test was performed. ^∗^p < 0.0332, ^∗∗^p < 0.0021, ^∗∗∗^p < 0.0002, and ^∗∗∗∗^<0.0001. **D)** Pie chart showing distribution of heavy chain sequences for donors VAIN1, VAIN2 and VAIN3. The number inside the circle represent the number of heavy chains analysed. The Pie slice size is proportional to the number of clonally related sequences and are colour coded based in clonal expansions described in **Supplementary Table 2**. The % on the outside of the Pie slice represents the overall % of sequences related to a clonal expansion. Graph showing the relative abundance of **(E)** V_H_ and **(F)** V_L_ gene usage for Spike-reactive mAbs isolated following infection, 2 doses of vaccine, 3 doses of vaccine or following BTI and IgG BCRs from SARS- CoV-2-naive individuals^36^. Statistical significance was determined by binomial test. *, P ≤ 0.05; **, P ≤ 0.01; ***, P ≤ 0.001; ****, P ≤ 0.0001. Blue stars are Vaccine vs Healthy, Red stars are BTI vs Healthy, Black are BTI vs Vaccine.

The degree of divergence from germline was also compared to a database of SARS-CoV-2 specific mAbs isolated from convalescent donors and individuals that had received 2 or 3 doses of COVID-19 vaccine^35^, as well as paired heavy and light chains of IgG B cell receptors from CD19+ B cells of healthy individuals^36^ (**Figure 2B and 2C**). Since the SARS-CoV-2 mAb database only included amino acid sequences for some mAbs, divergence from germline was determined at the amino acid level (which was previously shown to correlate well with nucleotide divergence^6^). BTI mAbs had a statistically higher amino acid mutation level (V_H_ 9.2% and V_L_ 6.2%) compared to mAbs isolated following infection only (V_H_ 4.2% and V_L_ 3.0%) and following two vaccine doses (V_H_ 5.3% and V_L_ 3.2%). However, there was no statistical difference in mutations levels between BTI mAbs and mAbs isolated following 3 vaccine doses (V_H_ 8.2% and V_L_ 5.6%) indicating an additional exposure to SARS-CoV-2 Spike in the form of infection or vaccination leads to increased somatic hypermutation. Non-Spike specific B cells were more highly mutated than BTI mAbs (V_H_ 10.9% and V_L_ 7.5%). Comparison of the CDRH3 length distribution with representative naive repertoires^37^ showed an enrichment in CDRH3 of lengths 20 amino acids (**Supplementary Figure 3D**) which is predominantly driven by a clonal expansion of a VH3-30 germline family from VAIN2 (**Supplementary Table 2**).

Sequence analysis identified clonally related sequences within all three donors (**Figure 2D** and **Supplementary Table 2**). Clonally expanded B cells represented 4%, 21% and 40% of all B cells from VAIN1, VAIN2 and VAIN3, respectively. Germline usage of BTI mAbs was also compared with non-Spike reactive mAbs and vaccine derived mAbs (**Figure 2E and 2F**).

As previously observed, there was an enrichment in VH3-53 and VH3-30/VH3-30-3 germline usage (**Figure 2E**)^5, 38–41^. mAbs utilizing these VH3-53 typically target the ACE2-binding site on RBD^5, 38–41^. VH3-30/VH3-30-3 encoded 20 RBD-specific mAbs with neutralizing activity (**Supplementary Table 3**). Enrichment in VH5-51 was seen for BTI mAbs compared to non-Spike mAbs from naïve donors but this was not seen for vaccine-derived mAbs. When comparing between vaccine-derived mAbs and BTI mAbs, enrichments in VH3-9, VH3-66 and VH3-13 were seen for vaccine-derived mAbs but not for BTI mAbs. When considering the light chain, enrichment in gene usage was seen for VK1-39 (22/119), VK1-5 (12/119) and VK1-33 (18/119) and enrichment of these germlines was greater than observed for vaccine-derived mAbs. Overall, there continues to be a diverse repertoire of SARS-CoV-2 specific mAbs present following infection in vaccinated individuals.

### mAbs isolated following BTI have broad neutralization against omicron sub-lineages

A selection of 67 neutralizing antibodies from the three donors were expressed and purified on a large scale for further characterization of neutralization breadth, potency and epitope specificity. Neutralization of purified mAbs was measured against a panel of viral particles pseudotyped with different SARS-CoV-2 variant Spikes, including WT, Delta, Beta, BA.1, BA.2 and BA.4/5 (**Supplementary Table 4**). mAbs with potent activity against all six viruses tested were identified in all three donors (**Figure 3A**). The neutralization potency of mAbs against WT and BTI variant correlated well for both delta BTI and BA.1 BTI mAbs (**Supplementary Figure 4**). When considering the geometric mean IC_50_ for mAbs isolated following VAIN1 and VAIN2 (delta infection) and VAIN3 (BA.1 infection), a different pattern of potencies was observed (**Figure 3A**). Whereas WT and delta were most potently neutralized by mAbs from the delta-infected donors, BA.1 and beta were most potently neutralized by mAbs from the BA.1-infected donor. BA.1 and beta share common mutations in RBD (K417N, E484K, N501Y) which could explain the high level of cross-reactivity of mAbs from VAIN3 with the beta variant.

**Figure 3:**
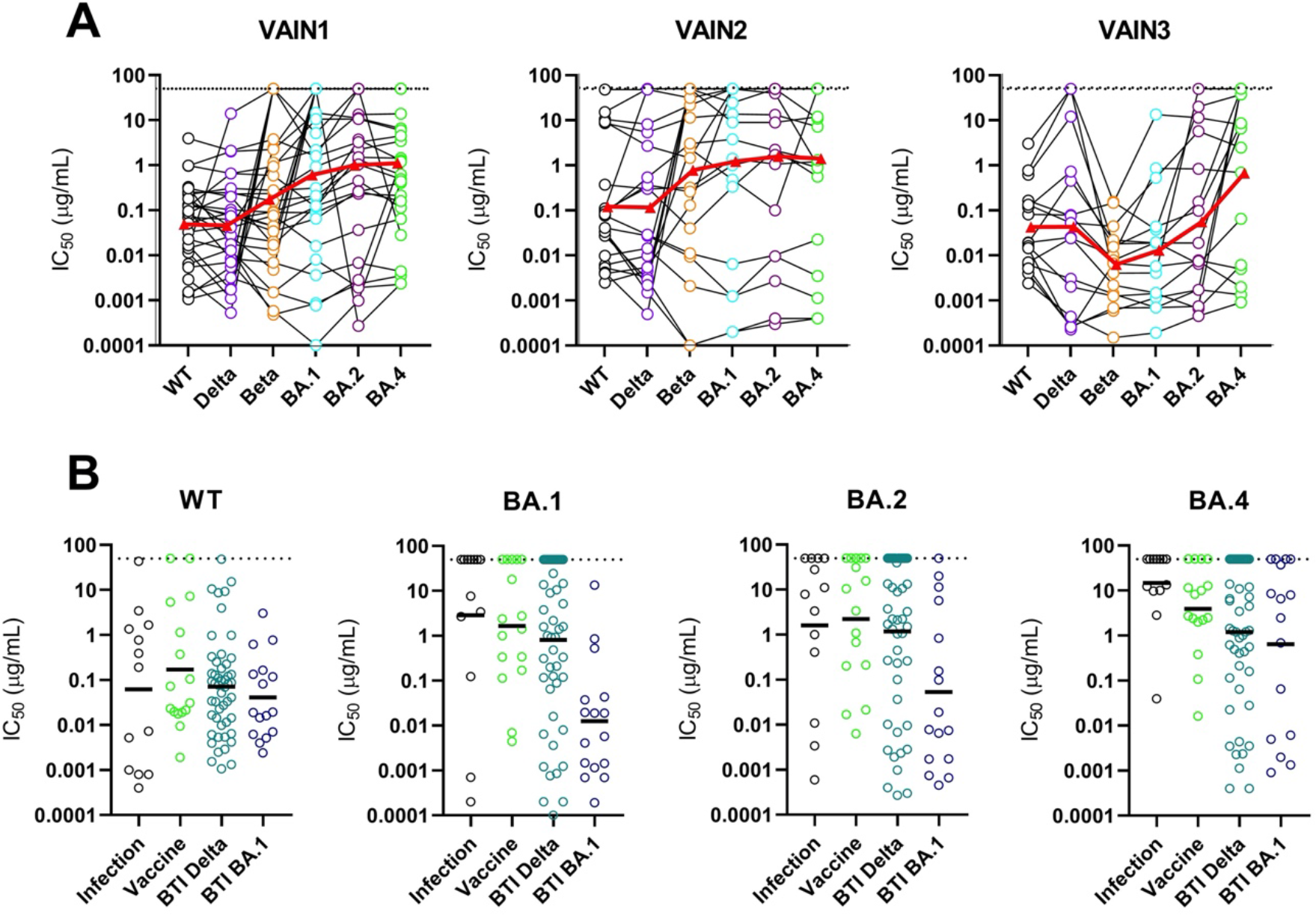
Neutralization breadth and potency against omicron sub-lineages. A) Neutralization breadth and potency of BTI mAbs against Wuhan-1, delta, beta, BA.1, BA.2 and BA.4 from VAIN1, VAIN2 and VAIN3. Data for each mAb is linked. Red triangle and linking line show the geometric mean IC_50_ against each variant. Dotted line represents the highest concentration of antibody tested. **B)** Comparison of neutralization breadth and potency of BTI mAbs with mAbs isolated from convalescent donors (Infection)^5^ and an AZD1222 vaccinated donor^6^ against omicron sub-lineages (BA.1, BA.2 and BA.4/5). Horizontal line represents the geometric mean IC_50_ against each mAb origin.

The neutralization breadth of mAbs isolated following BTI was compared to that of mAbs isolated from convalescent donors early in the pandemic (March – May 2020)^5^ and an mAbs isolated following 2-doses of AZD1222^6^. Analysis was focused on omicron sub-lineages BA.1, BA.2 and BA.4/5 (**Figure 3B**). The geometric mean IC_50_s against WT pseudotyped virus were most similar between the three mAb groups, whereas neutralization of the omicron sub- lineages showed larger differences. The lowest neutralization potencies were observed by infection and vaccine mAbs against BA.1, BA.2 and BA.4/5. mAbs isolated following delta BTI had similar GMT against BA.1, BA.2 and BA.4/5 whereas mAbs isolated following BA.1 BTI were more potent at neutralizing BA.1 compared to BA.2 and BA.4/5. The greater neutralization breadth of BTI mAbs is consistent with higher divergence from germline sequence.^6, 23, 42–44^ Interestingly, some of the mAbs isolated from the convalescent and AZD1222 vaccinated donors showed potent cross neutralization against all omicron sub- lineages^45^ despite having only experienced the WT Spike. Overall, mAbs with potent cross-neutralization were identified against antigenically omicron sub-lineages.

### RBD-specific mAbs form five competition groups

To understand more about the epitopes targeting on RBD, we performed Spike competition ELISAs between neutralizing antibodies with known RBD specificity that had been isolated from convalescent or vaccinated donors^5, 6^ (**Figure 4A-C** and **Supplementary Figure 5A**). Furthermore, to gain insight into mechanisms of neutralization, the ability of mAbs to inhibit binding of soluble Spike to HeLa-ACE2 cells was measured by flow cytometry^5^ (**Figure 4D**). mAbs with high inhibition levels directly block ACE2 binding through binding to the receptor binding motif (RBM)^5, 6^. The RBD-specific mAbs formed five competition groups (**Supplementary Figure 5A**), four of which had been observed previously^5, 6^. The distribution of RBD-specific mAbs between competition groups differed between the three donors, with VAIN1 and VAIN3 having the highest frequency of competition Group 4 and VAIN2 having the highest frequency of Group 3 (**Figure 4A**). Our previous studies isolating mAbs following infection^5^ or vaccination^6^ had shown a dominance of Group 3 and Group 4 RBD-specific mAbs, respectively. Interpretation of the biological significance of the differences in epitope immunodominance is limited due to the small number of mAbs studied.

**Figure 4:**
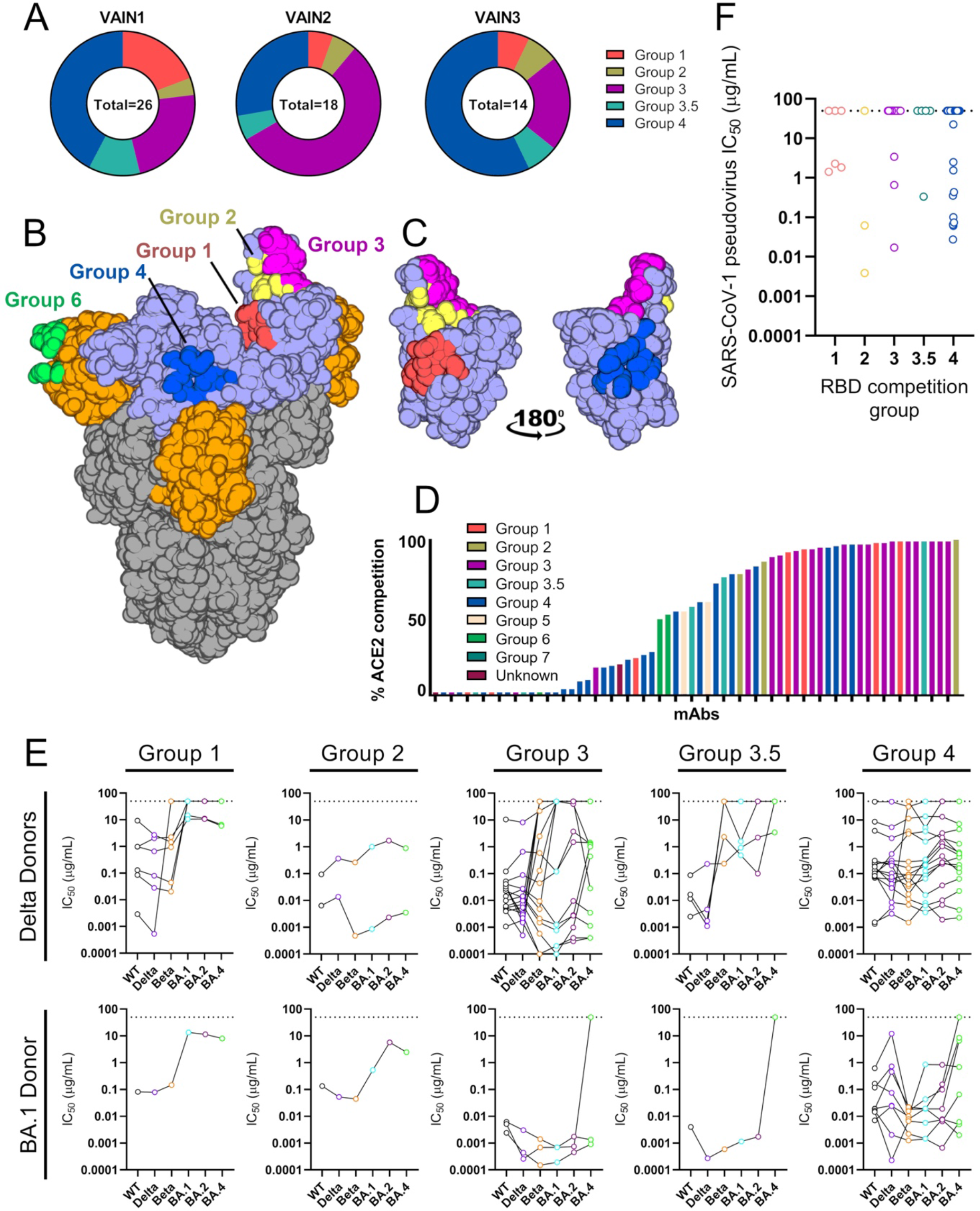
RBD mAb characterisation. A) Pie chart showing distribution of RBD-specific mAbs between competition groups for VAIN1, VAIN2 and VAIN3. **B)** Surface representation of SARS-CoV-2 WT spike (pdb:6XM0) showing epitopes of previously characterised competition groups as coloured surfaces^6^. RBD and NTD are indicated by light blue and orange, respectively. **C)** Surface representation of RBD domain in the up conformation showing location and proximity of group 1 (red), group 2 (yellow), group 3 (magenta) and group 4 (blue). Structures were generated in Pymol. **D)** Ability of RBD-specific neutralizing antibodies to inhibit the interaction between cell surface ACE2 and soluble SARS-CoV-2 Spike. mAbs (at 600 nM) were pre-incubated with fluorescently labelled Spike before addition to HeLa-ACE2 cells. The percentage reduction in mean fluorescence intensity is reported. Experiments were performed in duplicate. Bars are colour coded based on their competition group. **E)** Neutralization breadth and potency of RBD-specific mAbs within the different RBD competition groups. mAbs are separated by the infecting VOC. Data for each mAb is linked. Dotted line represents the highest concentration of antibody tested. **F)** Neutralization potency of RBD-specific mAbs against SARS-CoV-1. Data is presented by RBD competition group.

The majority of Group 1 mAbs, which bind an epitope distal to RBM (**Figure 4B and 4C**), showed neutralization activity against WT, delta and beta VOCS, but had greatly reduced or limited neutralization activity against the omicron sub-lineages (**Figure 4E**). This was true for mAbs isolated following both delta and BA.1 BTI. Group 2 mAbs, characterized by their ability to compete with both Group 1 and Group 3 mAbs, showed strong ACE2 competition (**Figure 4D**) as well as cross-neutralization of VOCs. Group 3 mAbs were enriched with VH3-53/3-66 germline usage (11/18) (**Supplementary Table 3**) which have been shown to bind the ACE2 receptor binding motif (RBM) on RBD^39–41^. Indeed, the majority of Group 3 mAbs showed >90% inhibition of ACE2 binding (**Figure 4D**). Interestingly, several mAbs that competed strongly with Group 3 mAbs showed very little inhibition of ACE2 binding suggesting a wide Spike footprint for this competition group and differing angles of approach. VH3-53/3-66 using mAbs showed broad and potent neutralization of omicron sub-lineages reaching IC_50_ <0.001 μg/mL (**Figure 4E**). However, Group 3 VH3-30 using mAbs isolated following delta BTI had limited neutralization breadth and only neutralized WT and delta VOCs (**Supplementary Table 3**).

mAbs within Group 4 competed with mAbs known to bind distal to the RBM and able to bind the RBD in its closed conformation (**Figure 4B**). These mAbs showed broad cross-neutralization across VOCs (**Figure 4E**) but the overall neutralization potency was reduced against WT and delta compared to the Group 3 mAbs with IC_50_ in the 0.001 – 8.65 μg/mL (geometric mean 0.11 μg/mL) and 0.0001 – 12.0 μg/mL (geometric mean 0.11 μg/mL) range for WT and delta, respectively (**Supplementary Figure 6**). There was an enrichment in VH5- 51 germline gene usage (6/24) (**Supplementary Table 3**). A range of ACE2 inhibitions were observed, indicating the large epitope footprint of this competition group (**Figure 4D**). An additional competition group (Group 3.5) was identified compared to our previous studies. Group 3.5 mAbs competed with both Group 3 and Group 4 mAbs (**Figure 4C**) and whilst potently neutralizing WT and delta, showed limited neutralization of omicron sub-lineages, in particular BA.4 (**Figure 4E**).

To determine whether the epitopes of the RBD-specific mAbs isolated following BTI are conserved on other betacoronaviruses, we next measured neutralization activity against SARS-CoV-1 pseudotyped virus (**Figure 4F**). Cross-neutralization of SARS-CoV-1 was observed for mAbs belonging to all five RBD competition groups, with a particular abundance within competition Group 4 (11/23). Group 2 mAb VAIN3O_12, isolated following BA.1 BTI was most potent, neutralizing SARS-CoV-1 with an IC_50_ of 0.0039 μg/mL.

Overall, RBD mAbs competed with previously isolated RBD-specific mAbs suggesting new RBD epitopes are not being targeted. However, the increased cross-competition between competition groups suggests a larger collective RBD footprint for neutralizing antibodies. Further structural characterisation is required to understand how VOC BTI influences specificity of RBD mAbs at the molecular level.

### Binding of NTD mAbs to VOCs does not correlate with neutralization activity

Competition for Spike binding between NTD-specific neutralizing antibodies with known specificity was used to determine the epitopes targeted by the seven NTD-specific neutralizing antibodies isolated^5^. We have previously identified three NTD-specific mAb competition groups^5, 6^ and structural characterization of mAb P008_56 from Group 6 revealed binding to NTD adjacent to the ß-sandwich fold^46^. The NTD-specific mAbs isolated following BTI formed three competition groups (**Supplementary Figure 5B**). Groups 5 and 6 were identified previously, but an additional group that did not compete with previously isolated NTD mAbs was also identified (designated NTD unknown). Group 5 mAbs VAIN2D_36 and VAIN2D_16 had poor cross-neutralization of VOCs (**Figure 5A**) and despite being isolated from a delta-infected donor, were unable to neutralize delta. Both Group 5 mAbs utilised the VH4-34 germline but were not clonally related. Whilst Group 6 mAbs showed greater cross-neutralization compared to Group 5 mAbs, none were able to neutralize all six VOCs (**Figure 5A**) but they were able to neutralize the variant the donor was infected with. The most broad and potent Group 6 mAb was VAIN1D_06 which neutralized all VOCs with IC_50_ < 0.55 μg/mL except BA.1. mAb VAIN1WT_13 from the non-competing group (NTD unknown) neutralized all 6 variants with IC_50_ between 0.033 – 13.9 μg/mL with the lowest neutralization potency against omicron sub-lineages.

**Figure 5:**
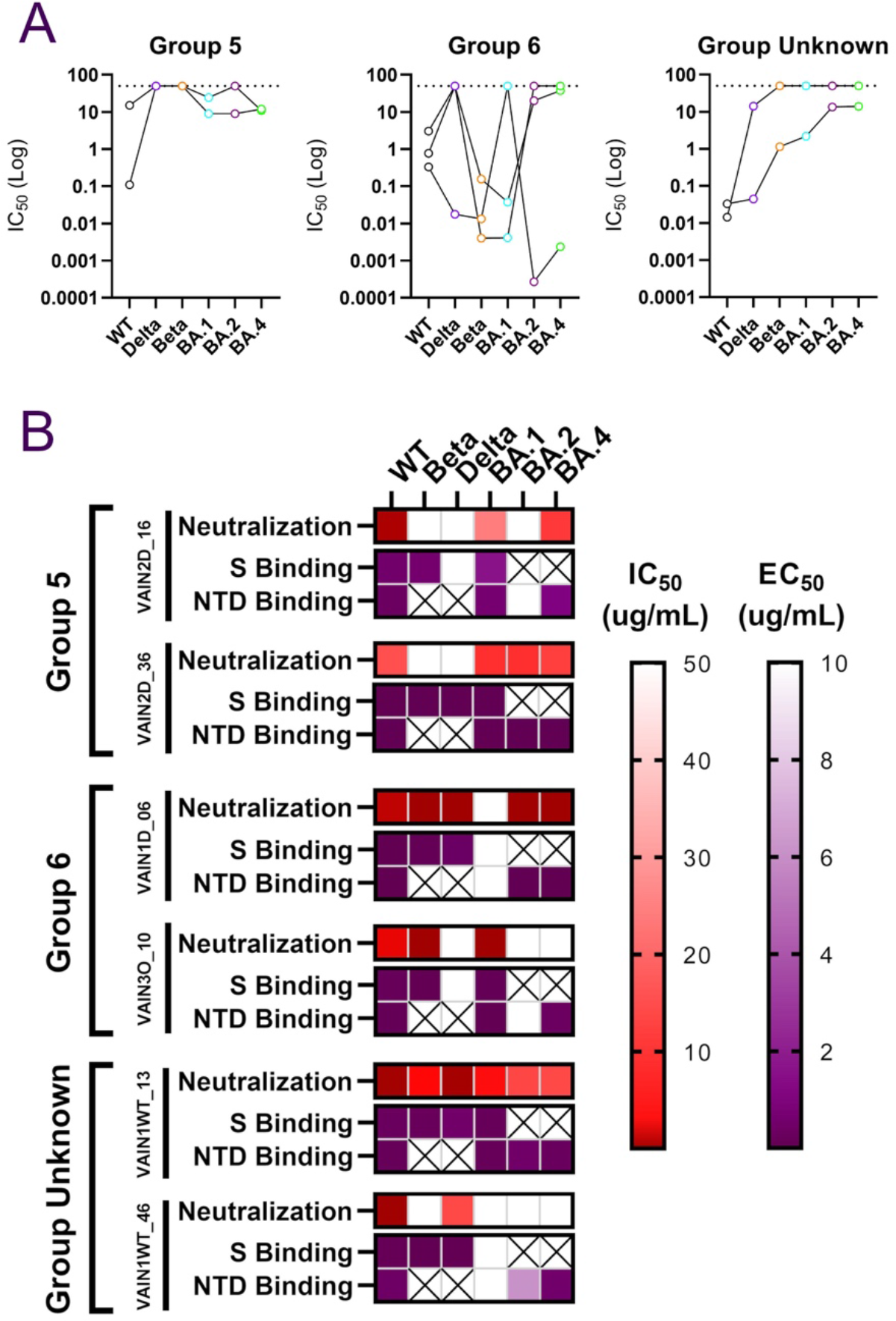
NTD mAb characterisation. A) Neutralization breadth and potency of NTD-specific mAbs within the different NTD competition groups. Data for each mAb is linked. Dotted line represents the highest concentration of antibody tested. **B)** Comparison between neutralization activity (IC_50_) and binding to Spike or NTD (EC_50_) by ELISA for NTD-specific mAbs. IC_50_ and EC_50_ values are shown as a heat map for each NTD-specific mAb with the level of binding shown in the key. A cross indicates that the Spike or NTD antigen for that variant was not tested.

To determine whether NTD mAbs lacking neutralization ability against a VOC was due to an inability to bind the NTD, ELISA assays were performed using recombinant Spike (WT, delta, beta, and BA.1) and recombinant NTD (WT, BA.1, BA.2 and BA.4) antigens (**Figure 5B**). Whilst binding and neutralization were consistent for VAIN1D_06 and VAIN1WT_13, binding did not always lead to neutralization for other NTD-specific mAbs. For example, Group 5 mAbs VAIN2D_16 and VAIN2D_36 bound well to recombinant beta Spike but did not neutralize beta pseudovirs. Furthermore, mAb VAIN1WT_46 bound to BA.2 and BA.4 NTD but did not neutralize the corresponding viral particles. This disconnect between NTD binding and neutralization was also observed by Wang *et al*.^47^ Mechanisms of NTD-specific mAb neutralization are not fully understood. However, the high mutation level in this region suggests NTD is under strong selective pressure from the host’s humoral immune response. McCallum *et al* demonstrate that some mAbs targeting the NTD supersite prevent SARS-CoV- 2 Spike mediated cell-cell fusion,^13^ whilst Cerutti *et al* showed that NTD mAbs use a restricted angle of approach to facilitate neutralization^48^. It is possible that the mutations, and/or insertions and deletions, within NTD encoded by different VOCs may alter the angle of approach which in turn reduces neutralization capability. Whether cross-binding but non-neutralizing NTD-specific mAbs can facilitate effector functions through their Fc receptors needs to be investigated further^49, 50^.

### XBB, XBB.1.5, BA.2.75.2 and BQ.1.1 show greater antigenic divergence

SARS-CoV-2 Spike continues to acquire mutations and since the omicron waves (including BA.1, BA.2, BA.4 and BA.5) new VOCs that have emerged include BA.2.75.2 (evolved from BA.2), XBB and XBB.1.5 (a recombinant of two BA.2 lineages, BA.2.75 and BJ.1) and BQ.1.1 (evolved from BA.5) (**Supplementary Table 4**). Despite these variants being on divergent evolutionary courses, they share convergent mutations in RBD. Additional mutations in RBD compared to BA.1 include R326T and N460K in BA.2.75.2, XBB/XBB.1.5 and BQ.1.1, G446S and F486S in BA.2.75.2 and XBB/XBB.1.5, and K444T in BQ.1.1. A panel of mAbs were selected, based on their neutralization activity against omicron sub-lineages, to gain insight into whether BTI after two vaccine doses could elicit antibodies capable of neutralizing these new variants (**Figure 6B-C**). Neutralization potencies were compared to the neutralization activity in sera from the three donors (**Supplementary Figure 7**) as well as with a larger group of double vaccinated individuals experiencing a delta BTI (**Figure 6A**).

**Figure 6:**
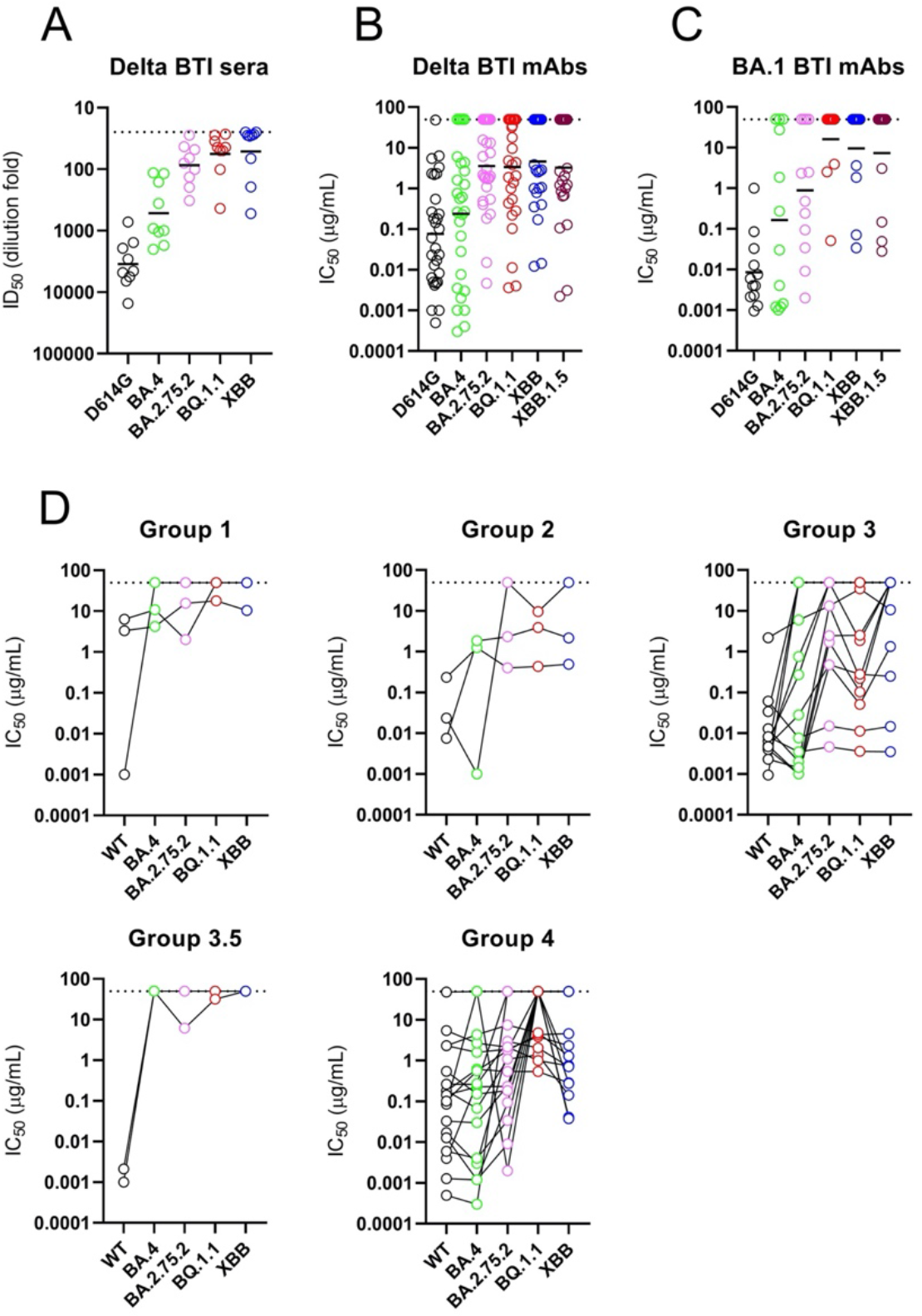
mAb neutralization against more recent VOCs including BA.2.75.2, XBB/XBB.1.5 and BQ.1.1. Neutralization by **A)** plasma from individuals vaccinated with 2 doses of BNT162b2 and were subsequently delta-infected (ID_50_), **B)** mAbs from delta BTI donors (IC_50_), and **C)** mAbs from BA.1 BTI donor (IC_50_). Sera was collected 15-35 days post infection. Additional plasma samples from double vaccinated and BA.1 infected indiviudals were not available. Horizontal line represents geometric mean IC_50_ (for mAbs) or geometric mean titres (plasma). **D)** Neutralization breadth and potency broken down by RBD competition group. Data for each mAb is linked. Dotted line represents the highest concentration of mAb or lowest dilution of plasma tested.

Whereas the sera from donors VAIN1, VAIN2 and VAIN3 had shown broad cross-neutralization of omicron sub-lineages (**Supplementary Figure 2**), there was a reduction in neutralization of BA.2.75.2, XBB, XBB.1.5 and BQ.1.1 (**Supplementary Figure 7**). This pattern of neutralization was observed in the larger panel of sera tested (**Figure 6A**) as well as by the isolated mAbs (**Figure 6B and 6C**). Whilst many of the BTI mAbs had retained some level of neutralization activity against the omicron sub-lineages, many of the mAbs tested lost neutralization activity against all four VOCs. However, RBD-specific mAbs with cross-neutralizing activity against all variants were still identified and importantly these belonged to multiple RBD competition groups (including Group 2, Group 3 and Group 4) (**Figure 6D**) indicating that the additional Spike mutations did not lead to complete disruption of all RBD neutralizing epitopes. Group 3 mAbs, VAIN2D_12 and VAIN2D_17, were most potent, neutralizing all VOCs with IC_50_ <0.01 μg/mL (**Supplementary Table 3**). Other cross-neutralizing RBD-specific antibodies were less potent, only reaching IC_50_s between 0.1 μg/mL and 10 μg/mL. Whether mAbs with cross-neutralizing activity could undergo further mutation to enhance neutralization potency would be of interest for optimization of mAbs for therapy against diverse VOCs.

Overall, despite broad neutralization of BA.1, BA.2 and BA.4, the convergent RBD mutations in BA.2.75.2, XBB/XBB.1.5 and BQ.1.1 lead to extensive immune evasion to mAbs generated following delta and BA.1 BTI. Several potent cross-neutralizing mAbs were identified and additional structural studies would provide important insights into how these mAbs tolerate these additional RBD mutations.

## Discussion

Studies conducted by us and others using convalescent sera or plasma have shown that a delta or BA.1 infection following COVID-19 vaccination can broaden the neutralization activity against omicron sub-lineages^19, 20, 22, 51, 52^. Through isolation of mAbs from BNT162b2 double vaccinated individuals that were subsequently delta or BA.1-infected, we showed that this increase in neutralization breadth is due to the presence of mAbs with potent cross-neutralizing activity. Despite using antigen baits specific for the vaccine and the infecting variant, we observed similar levels of WT and VOC specific B cells, and did not identify mAbs that were specific for the infecting variant. Combined with the observation that BTI mAbs had a higher level of somatic hypermutation compared to vaccine and infection-only elicited mAbs, we infer that delta or BA.1 infection in vaccinated individuals predominantly resulted in re- activation and maturation of B cells generated through previous COVID-19 vaccination rather than a *de novo* response specific to the VOC Spike, consistent with several other recent studies on breakthrough infection^23, 53–55^.

With the reactivation of existing B cells, it might be expected that immune imprinting from prior COVID-19 vaccinations based on Wuhan-1 might limit neutralization breadth of mAbs in a manner similar to that observed following influenza re-exposure^26–28^. However, the continued maturation upon re-activation of B cells leads to mAbs with increased neutralization breadth. This observation is supported by research showing that wider SARS-CoV-2 neutralization breadth is associated with increased somatic hypermutation^6, 23, 42–44^. The three donors studied here had received two vaccine doses prior to infection with an antigenically distinct Spike (either delta or BA.1). A third vaccine dose based on the Wuhan-1 strain has also been shown to increase neutralization breadth within polyclonal sera/plamsa^19, 56, 57^ and mAbs isolated from such individuals also show continued maturation and increased neutralization breadth and potency against VOCs, in particular BA.1^29, 30, 58^. Combined, these findings show that a third antigenic stimulation, independent of the Spike variant, can increase neutralization breadth. However, studies examining the impact of a fourth antigenic stimulation show a more modest increase in neutralization breadth and potency of isolated mAbs^54^.

This study has implications for variant based vaccine boosters. COVID-19 vaccine boosters are important for maintaining circulating levels of antibodies as well as providing broadened protection against newly emerging variants. Whilst the goal of variant based vaccine boosters is to match circulating strains dominant in the human population, these findings suggest that exposure to an antigenically distinct Spike (either delta of BA.1) can provide broad protection through generating mAbs with cross-neutralizing activity instead of eliciting a *de novo* response specific for the variant. Indeed, bivalent vaccine boosters based upon the BA.1 or BA.4/5 Spike antigens are now being used, and are effective at generating broad neutralization against omicron sub-lineages similar to monovalent boosters^59, 60^ and are effective at preventing severe disease following BA.4.6, BA.5, BQ.1, and BQ.1.1 infections^61^. Many BTI mAbs could neutralize variants of concern which have diverged independently from the ancestral Wuhan-1 strain and previous studies using a variety of immune sera have highlighted their antigenic distance^62, 63^. This cross-neutralization highlights that despite large variation in Spike, several conserved neutralizing epitopes exist on RBD, and to a lesser extent NTD. This is further exemplified by identification of RBD-specific mAbs from all five competition groups that have neutralization activity against SARS-CoV-1. Interestingly, mAbs isolated following BA.1 BTI had greater cross-neutralization of beta compared to mAbs isolated following delta infection. BA.1 and beta share common mutations in RBD (K417N, E484K, N501Y) and mAbs directed against these mutated epitopes could explain the high level of cross-reactivity of mAbs from VAIN3 with beta.

The numbers of mAbs isolated are too small to draw strong conclusions regarding differences in epitope immunodominance upon different variant exposure. However, it is clear that neutralization activity converges on similar Spike epitopes. Since the delta and BA.1 infection waves, SARS-CoV-2 has continued to mutate. Many of the BTI mAbs isolated here lost or had greatly reduced neutralization activity against currently circulating VOCs (i.e. early 2023), including BA.2.75.2, XBB/XBB.1.5 and BQ.1.1. This pattern was also observed in sera/plasma from BTI individuals and has been reported by several other groups worldwide^64–67^. This suggests that mAbs generated during the large delta and BA.1 infection waves between June 2021 to March 2022 may have acted as selective pressures in driving immune escape of these VOCs, in particular selecting mutations within RBD. Indeed, BA.2.75.2 was highly prevalent in India following a large delta wave^68^. BA.2.75.2, XBB/XBB.1.5 and BQ.1.1. converge in their RBD mutational profile. BA.2.75.2, XBB/XBB.1.5 and BQ.1.1. share common mutations including R346T (within the competition Group 4 epitope) and N460K (within the competition Group 1 epitope), and XBB and BA.2.75.2 also share G446S and F486S mutations (within competition Group 3 epitope). Wang *et al* demonstrated that introduction of R346T, K444T or N460K into BA.4/5 and R346T, V445P or N460K into BA.2 were responsible for reduction in neutralization by many RBD-specific mAbs^64^. However, the identification of mAbs belonging to several RBD competition groups that had neutralization activity against all VOCs tested suggests that multiple additional Spike mutations would be required across RBD to generate complete immune evasion of the antibody response following BTI. The mAb recall response was diverse in gene usage despite multiple clonal expansions being observed. Maintaining a diverse response would not only limit immune escape through selection of Spike mutations but may also represent a wide pool of B cells that could be re-activated by a diverse range of antigenically distinct Spike variants. Further studies characterizing the antibody-Spike interaction at the molecular level would provide information on how these mAbs retain cross-neutralizing activity despite high-levels of Spike mutations and may help to predict future Spike escape variants.

Encouragingly, RBD-specific mAbs with cross-neutralizing activity against the most recent VOCs (BA.2.75.2, XBB/XBB.1.5 and BQ.1.1) were found within four RBD competition groups. These less frequent mAbs represent potential candidates for the next generation of antibody-based therapeutics against SARS-CoV-2 and other betacoronaviruses. Although these mAbs represent a minor component of the mAb response, understanding how to selectively boost these responses could aid in preparedness against new SARS-CoV-2 variants as they arise. Overall, infection with a VOC following two COVID-19 vaccine doses shapes the antibody response by re-activating and maturing existing Spike specific B cells to produce mAbs with broad neutralization activity.

## Methods

### Ethics and samples

This study used human samples collected with written consent as part of a study entitled “Antibody responses following COVID-19 vaccination”. Ethical approval was obtained from the King’s College London Infectious Diseases Biobank (IBD) (KDJF-110121) under the terms of the IDB’s ethics permission (REC reference: 19/SC/0232) granted by the South Central Hampshire B Research Ethics Committee in 2019, and London Bridge Research Ethics Committee (Reference: REC14/LO/1699). Collection of surplus serum samples at St Thomas Hospital, London, was approved by South Central-Hampshire B REC (20/SC/0310). SARS-CoV-2 cases were diagnosed by either reverse transcriptase PCR (RT-PCR) of respiratory samples at St Thomas’ Hospital, London, UK or by lateral flow testing. All participants were SARS-CoV-2 naïve prior to vaccination and infection. Participants VAIN1 and VAIN2 were infected during the UK Delta wave (11/8/21 and 23/8/21, respectively), and participant VAIN3 was infected during the UK BA.1 wave (18/12/21). Viral sequencing was not performed on these samples.

### Antigen-specific B cell sorting

Fluorescence-activated cell sorting of cryopreserved PBMCs was performed on a BD FACS Melody as previously described^5, 6^. Sorting baits with a Strep2A tag (SARS-CoV-2 Wuhan S1, Delta S1 and BA.1 S1) was pre-complexed with the StrepTactin fluorophore at a 1:1 molar ratio prior to addition to cells. PBMCs were stained with live/dead (fixable Aqua Dead, Thermofisher), anti-CD3-APC/Cy7 (Biolegend), anti-CD8-APC-Cy7 (Biolegend), anti- CD14-BV510 (Biolegend), anti-CD19-PerCP-Cy5.5 (Biolegend), anti-IgM-PE (Biolegend), anti-IgD-Pacific Blue (Biolegend) and anti-IgG-PeCy7 (BD) and S1-StrepTactin XT DY-649 (IBA life sciences, 2-1568-050) and S1-StrepTactin XT DY-488 (IBA life sciences, 2-1562- 050). Live CD3/CD8^-^CD14^-^CD19^+^IgM^-^IgD^-^IgG^+^S1^+^S1^+^ cells were sorted using a BD FACS Melody into individual wells containing RNase OUT (Invitrogen), First Strand SuperScript III buffer, DTT and H_2_O (Invitrogen) and RNA was converted into cDNA (SuperScript III Reverse Transcriptase, Invitrogen) using random hexamers (Bioline Reagents Ltd) following the manufacturer’s protocol.

### Full-length antibody cloning and expression

The human Ab variable regions of heavy and kappa/lambda chains were PCR amplified using previously described primers and PCR conditions^31, 32, 69^. PCR products were purified and cloned into human-IgG (Heavy, Kappa or Lambda) expression plasmids^32^ using the Gibson Assembly Master Mix (NEB) following the manufacturer’s protocol. Gibson assembly products were directly transfected into HEK-293T/17 cells and transformed under ampicillin selection. Ab supernatants were harvested 3 days after transfection and IgG expression and Spike-reactivity determined using ELISA. Ab variable regions of heavy-light chain pairs that generated Spike reactive IgG were sequenced by Sanger sequencing.

Antibody heavy and light plasmids were co-transfected at a 1:1 ratio into HEK-293F cells (Thermofisher) using PEI Max (1 mg/mL, Polysciences, Inc.) at a 3:1 ratio (PEI Max:DNA). Ab supernatants were harvested five days following transfection, filtered and purified using protein G affinity chromatography following the manufacturer’s protocol (GE Healthcare).

### Pseudovirus production

HEK293T/17 cells were seeded the day prior on 10 cm dishes at a density of 7x10^5^ cells/mL in DMEM with 10% FBS, 1% Pen-Strep. Cells were co-transfected using 90 µg PEI-Max (1 mg/mL, Polysciences) with 15 µg HIV-luciferase plasmid, 10 µg HIV 8.91 gag/pol plasmid^70^ and 5 µg SARS-CoV-2 Spike protein plasmid. Transfected cells were incubated for

72 h at 37°C and virus was harvested, sterile filtered and stored at -80°C until required. Mutations present in each variant Spike are shown in **Supplementary Table 4**.

### Neutralization assays

Serial dilutions of plasma or mAb in DMEM, supplemented with 10% fetal bovine serum (FBS) and 1% Pen/Strep, were incubated in a 96 well plate, with HIV-1 virus pseudotyped with SARS-CoV-2 wild-type or variant Spikes for 1 h at 37°C. HeLa cells stably expressing the human ACE2 receptor were then added at a density of 4x10^5^ cells/mL to all wells and incubated for 72 h at 37°C. Levels of infection was measured with the Bright-Glo luciferase kit (Promega) on a Victor X3 multilabel reader (Perkin Elmer). Duplicate measurements were used to calculate IC_50_ and ID_50_.

### ELISA (Spike, RBD, NTD, or S1)

96-well plates (Corning, 3690) were coated with Spike, S1, NTD or RBD at 3 μg/mL overnight at 4°C. The plates were washed (5 times with PBS/0.05% Tween-20, PBS-T), blocked with blocking buffer (5% skimmed milk in PBS-T) for 1 h at room temperature. Serial dilutions of mAb or supernatant in blocking buffer were added and incubated for 2 hr at room temperature. Plates were washed (5 times with PBS-T) and secondary antibody was added and incubated for 1 hr at room temperature. IgG was detected using Goat-anti-human-Fc-AP (alkaline phosphatase) (1:1,000) (Jackson: 109-055-098). Plates were washed (5 times with PBS-T) and developed with either AP substrate (Sigma) and read at 405 nm.

### Competition ELISA

F(ab’)_2_ of previously characterized mAbs were produced by IdeS digestion of IgG as described previously^5^. 96-well plates (Corning, 3690) were coated with WT Spike at 3 μg/mL overnight at 4°C. Plates were washed and blocked as described above. Serial dilutions (5- fold) of F(ab’)_2_, starting at 100-fold molar excess of the EC_80_ of Spike binding were added to the plate and incubated for 1 h at room temperature. Plates were washed (5x with PBS-T) and competing IgG was added at the EC80 of Spike binding and incubated for 1 h at room temperature. Plates were washed (5x with PBS-T) and Goat-anti-human-Fc-AP (alkaline phosphatase) (1:1,000) (Jackson: 109-055-098) was added and incubated for 1 h at room temperature. The plates were washed a final time (5x with PBS-T) and the plate was allowed to develop by addition of AP substrate (Sigma). Optical density at 405 nm was measured in 5 min intervals. Percentage competition was calculated using the equation below and competition group clusters were arranged by hand according to binding epitope.

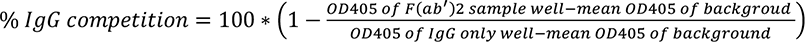

### ACE2 competition measured by flow cytometry

Fluorescent probe was prepared by mixing 3.5 molar excess of Streptavidin-APC (Thermofisher Scientific, S32362) with biotinylated SARS-CoV-2 spike and incubating for 1 h on ice. Purified mAb was mixed with APC conjugated Spike in a molar ratio of 4:1 in FACS buffer (2% FBS in PBS) on ice for 1h. HeLa-ACE2 cells were washed once with PBS and detached using 5 mM EDTA, PBS. Cells were washed and resuspended in FACs buffer before adding 5x10^5^ cells to each mAb-Spike complex. Cells were incubated on ice for 30 min. HeLa- ACE2 cells alone and with SARS-CoV-2 Spike only were used as background and positive controls, respectively. The geometric mean fluorescence of APC was measured from the gate of singlet cells. ACE2 binding inhibition was calculated with this equation:

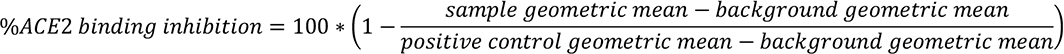

### Sequence analysis of Monoclonal antibodies

Heavy and light chain sequences of SARS-CoV-2 specific mAbs were examined using IMGT/V-quest (http://www.imgt.org/IMGT_vquest/vquest) to identify germline usage, percentage of SHM and CDR region lengths. 5 amino acids or 15 nucleotides were truncated from the start and end of the sequences to remove variation introduced from the use of a mixture of forward cloning primers. D’Agostino and Pearson tests were performed to determine normality. Based on the result, a Kruskal-Wallis test with Dunn’s multiple comparison post hoc test was performed. Two-sided binomial test was performed in excel. Significance defined as p < 0.0332 (*), 0.0021 (**), 0.0002 (***) and >0.0001 (****).

## Acknowledgements

We thank Philip Brouwer, Marit van Gils and Rogier Sanders for the Spike protein construct, Peter Cherepanov for S1 proteins from VOCs, Wendy Barclay for providing the beta, delta, BA.1, BA.2, BA.4/5, BA.2.75.2, XBB, XBB.1.5 and BQ.1.1 Spike plasmids and James Voss and Deli Huang for providing the Hela-ACE2 cells.

