## Supplemental Information for "Broad and potent neutralizing mAbs are elicited in vaccinated individuals following Delta/BA.1 breakthrough infection"

* These authors contributed equally

**Supplementary Figure 1: Sorting strategy for isolation of S1 reactive B cells following infection in vaccinated individuals.** Strategy to isolate SARS-CoV-2 S1 specific IgG+ B cells. Example sorting for donor VAIN1. Live CD3/CD8$^-$CD14$^-$CD19$^+$IgM$^-$IgD$^-$IgG$^+$S1$^+$S1$^+$ cells were sorted into individual wells. The heavy and light chains were reverse transcribed and amplified using nested PCR with gene specific primers[1-3].

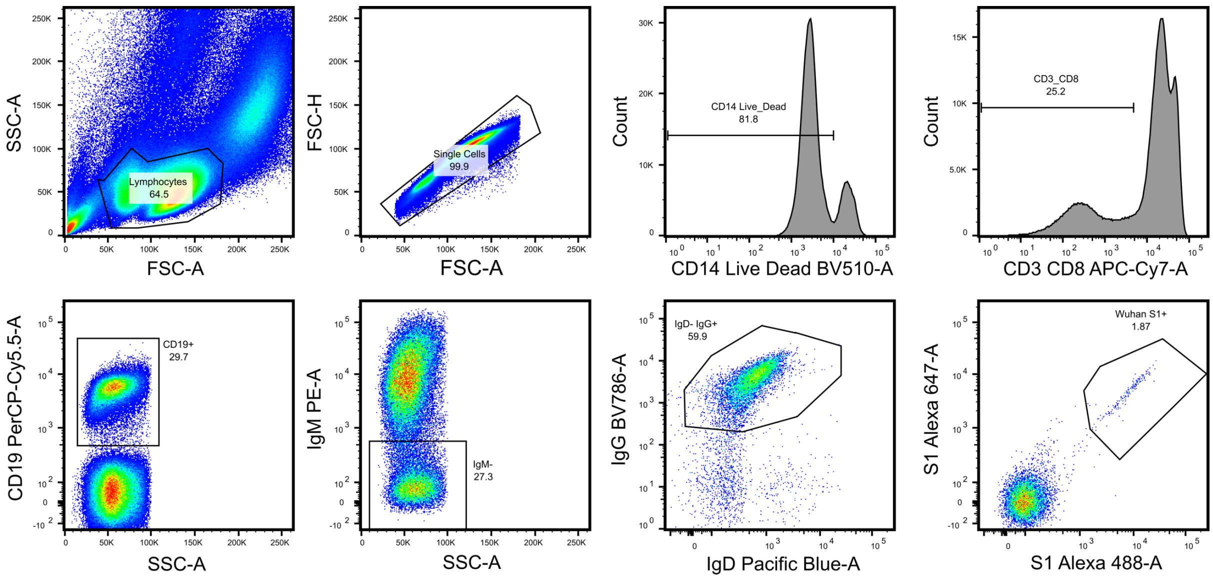


**Supplementary Figure 2: Plasma cross-neutralizing activity for donors VAIN1, VAIN2 and VAIN3.** Neutralization was tested using HIV-1 viral particles pseudotyped with Spike of Wuhan-1 (WT), beta, delta, BA.1, BA.2 and BA.4/5.

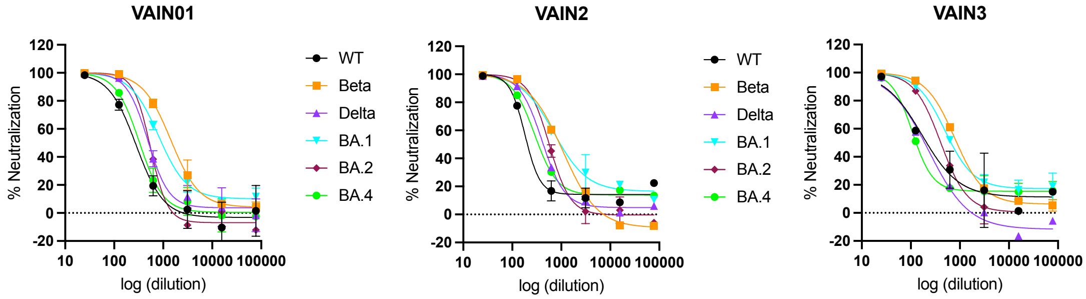

**Supplementary Figure 3: Comparison between mAbs isolated from VAIN1, VAIN2 and VAIN3. A)** Truncated violin plot comparing the level of nucleotide mutation from germline for $V_H$ and $V_L$ between mAbs isolated from VAIN1, VAIN2 and VAIN3. **B)** Truncated violin plot comparing the level of nucleotide mutation from germline for $V_H$ and $V_L$ between B cells selected using WT S1, Delta S1 or BA.1 S1. **C)** Plot comparing the level of nucleotide mutation between WT and VOC selected B cells for donors VAIN1, VAIN2 and VAIN3, respectively. D'Agostino and Pearson tests were performed to determine normality. Based on the result, a Kruskal-Wallis test with Dunn's multiple comparison post hoc test was performed. $^{*}p < 0.0332$, $^{**}p < 0.0021$, $^{***}p < 0.0002$, and $^{****}<0.0001$. **D)** Distribution of CDRH3 lengths for mAbs isolated following BTI and representative naive B cell repertoire[4]. Error bars represent the standard deviation between donors used in the analysis (n = 3 for BTI mAbs and n = 10 for naive repertoire). A bimodal distribution of CDRH3 length is observed for SARS-CoV-2 Spike reactive mAbs.

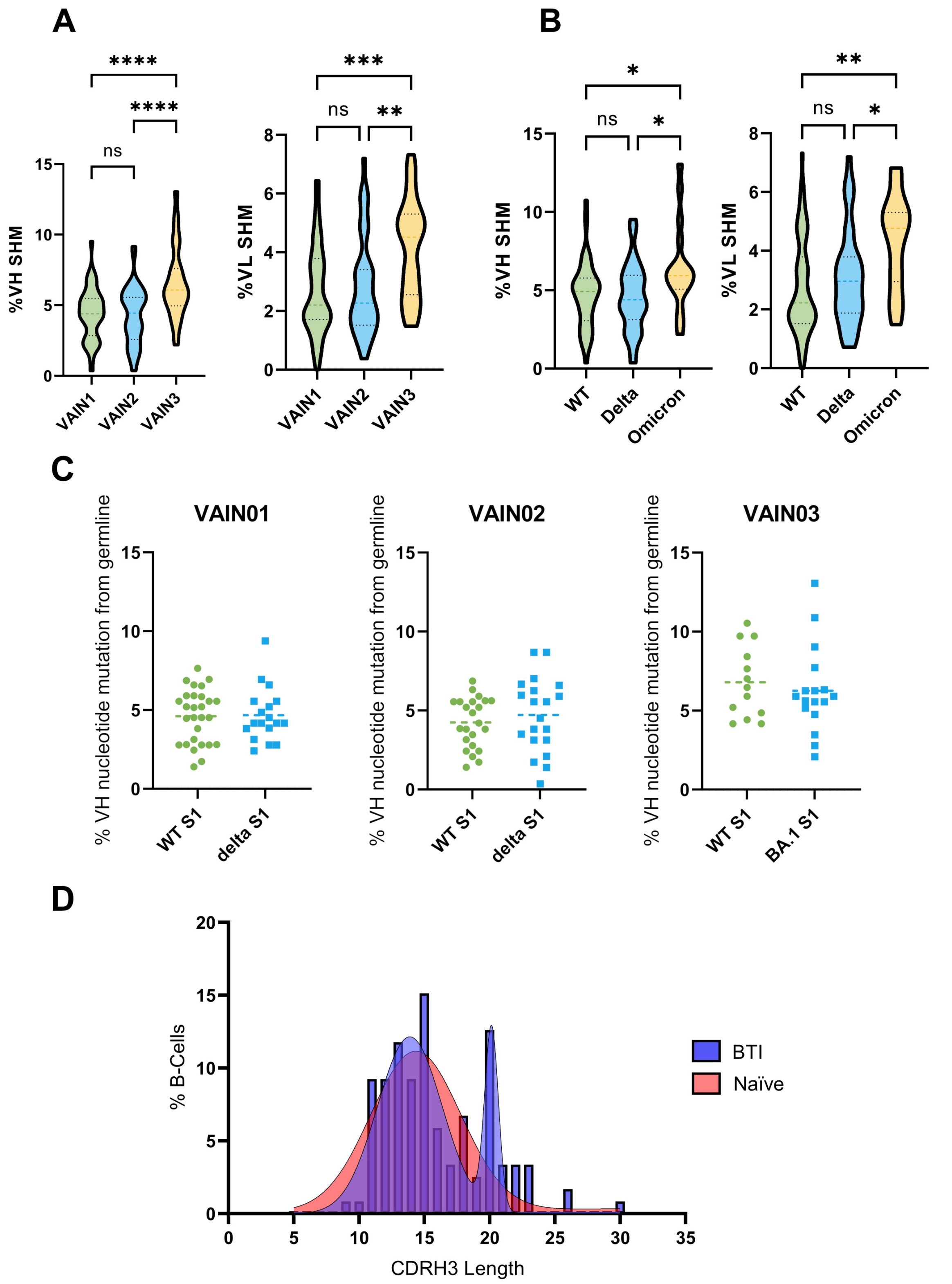
A
****
****
ns
15
10
5
0
%VH SHM
VAIN1
VAIN2
VAIN3
***
ns
**
8
6
4
2
0
%VL SHM
VAIN1
VAIN2
VAIN3
B
*
ns
*
15
10
5
0
%VH SHM
WT
Delta
Omicron
**
ns
*
8
6
4
2
0
%VL SHM
WT
Delta
Omicron
C
VAIN01
VAIN02
VAIN03
% VH nucleotide mutation from germline
WT S1
delta S1
WT S1
delta S1
WT S1
BA.1 S1
D
20
15
10
5
0
% B-Cells
0
5
10
15
20
25
30
35
CDRH3 Length
BTI
Naïve

**Supplementary Figure 4: Correlation between WT and BTI VOC neutralization potency.** Correlation between $IC_{50}$ values against WT and delta (VAIN1 and VAIN2 mAbs) or against WT and BA.1 (VAIN3 mAbs). Delta correlation is shown in purple and BA.1 correlation is shown in blue. (Spearman correlation, r. A linear regression was used to calculate the goodness of fit, $r^2$).

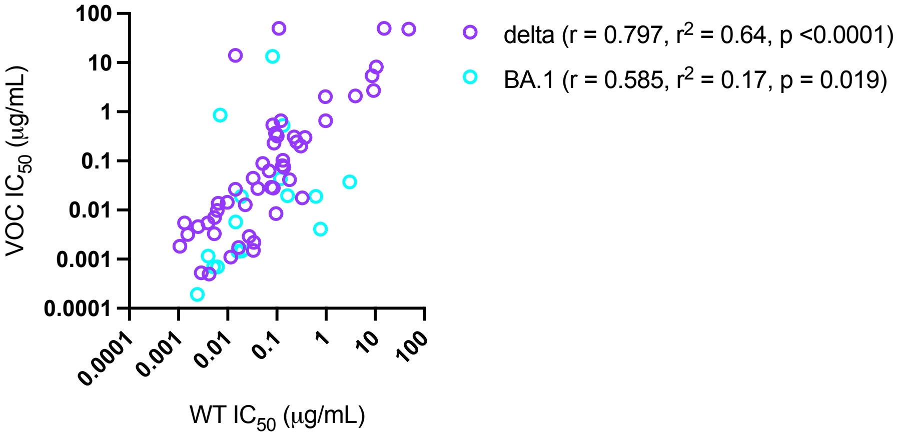

**Supplementary Figure 5: RBD- and NTD-specific mAbs form multiple competition groups.** Competition for **A)** RBD-specific mAbs and **B)** NTD-specific mAbs. Inhibition of IgG binding to SARS-CoV-2 Spike by F(ab)$_2$' fragments was measured. The percentage competition was calculated using the reduction in IgG binding in the presence of F(ab')$_2$ (at 100-molar excess of the IC$_{80}$) as a percentage of the maximum IgG binding in the absence of F(ab')$_2$. Competition groups clusters were arranged by hand according to binding epitopes. Experiments were performed in duplicate. Competition <25% is white. Grey boxes indicate competition not tested. Competition groups are colour-coded according to the key.

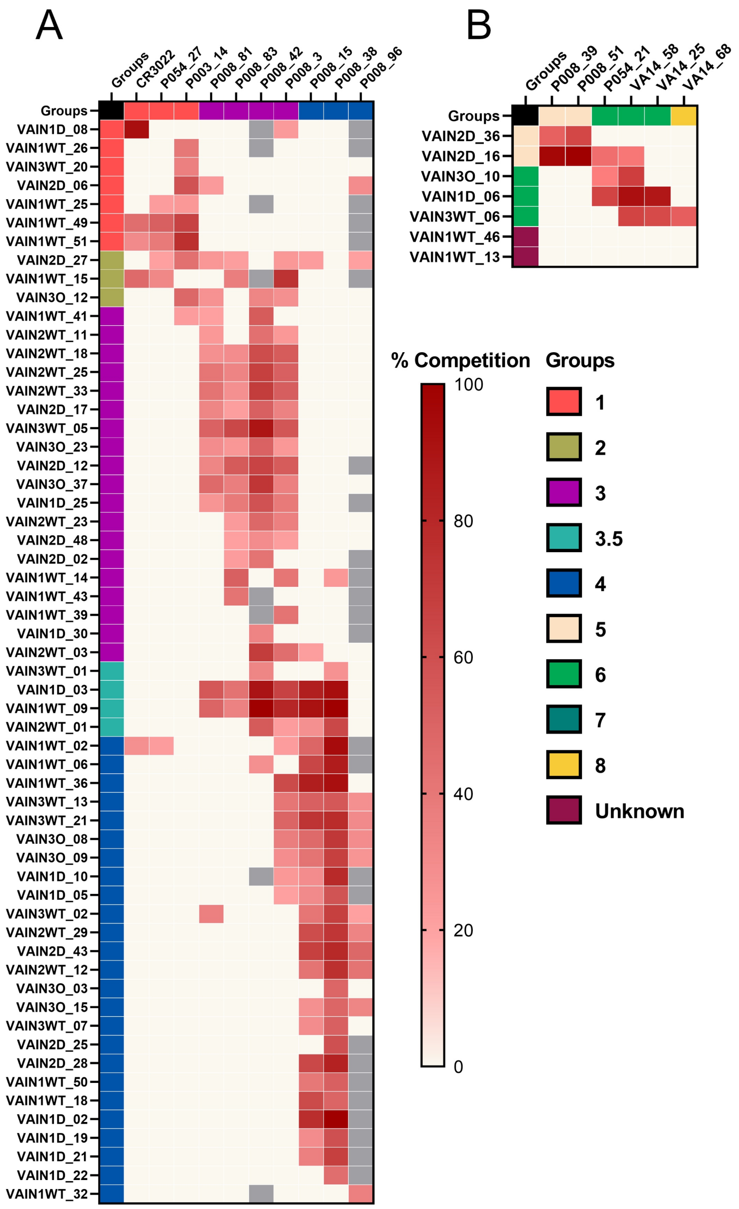

**Supplementary Figure 6: RBD-specific mAb neutralization geometric mean $IC_{50}$ against SARS-CoV-2 VOCs by competition group.** Dotted line represents the highest mAb concentration tested. The horizontal line shows the geometric mean $IC_{50}$ for each RBD competition group.

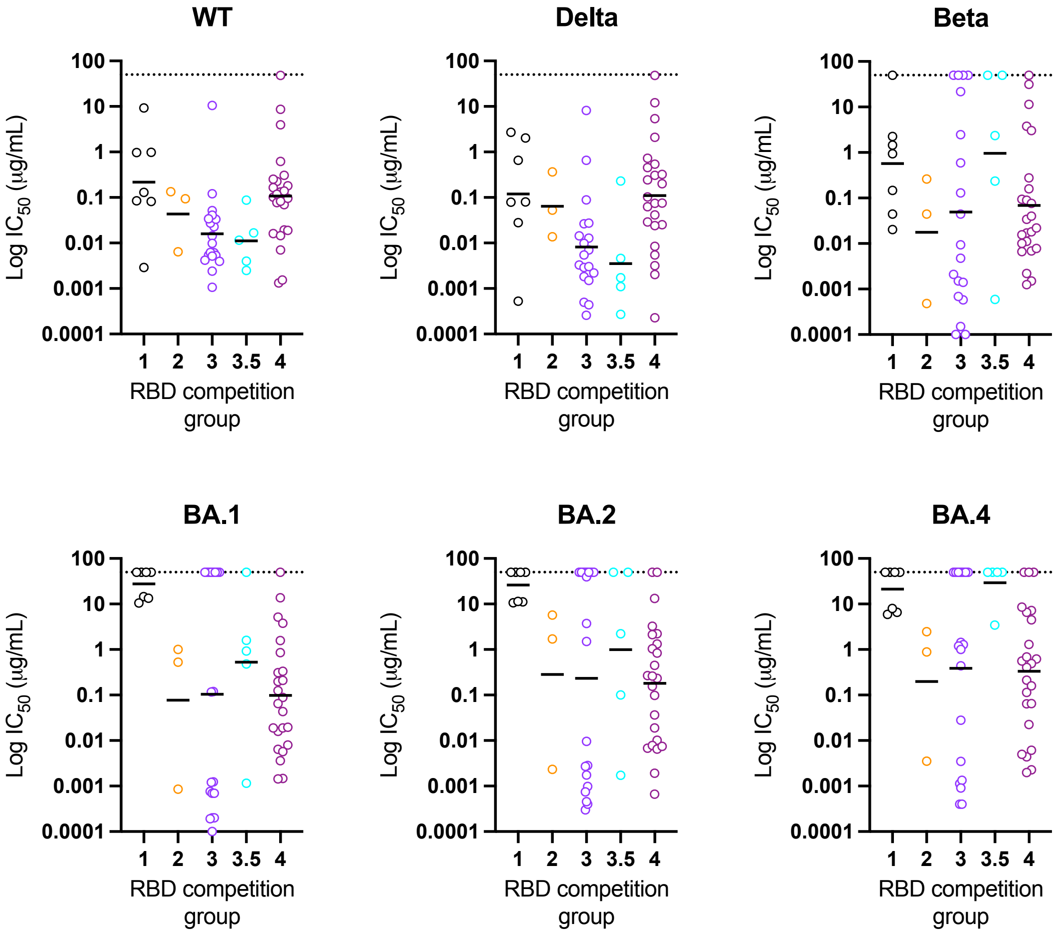


**Supplementary Figure 7: VAIN1, VAIN2 and VAIN3 plasma neutralization against BA.2.75.2, XBB and BQ.1.1.**

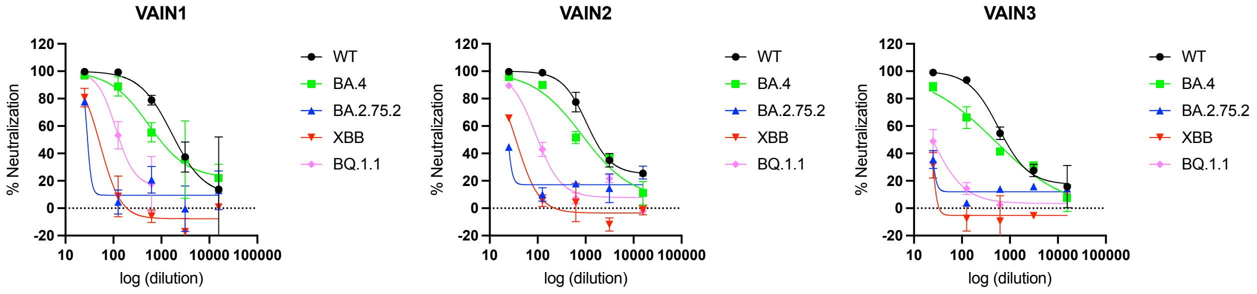

**Supplementary Table 1: VAIN1, VAIN2 and VAIN3 donor information**. Dates for vaccination and SARS-CoV-2 infection and the days between these events and blood donation.

| | VAIN1 | VAIN2 | VAIN3 |
|---|---|---|---|
| *Gender* | Female | Male | Female |
| *Ethnicity* | White | White | White |
| *Age* | na | 42 | 32 |
| *1st Vaccine (Vaccine type)* | 12/01/2021 (Pfizer mRNA:BNT162b2) | 21/01/2021 (Pfizer mRNA:BNT162b2 | 21/05/2021 (Pfizer mRNA:BNT162b2) |
| *2nd Vaccine (Vaccine type)* | 02/03/2021 (Pfizer mRNA:BNT162b2 | 03/03/2021 (Pfizer mRNA:BNT162b2) | 01/07/2021 (Pfizer mRNA:BNT162b2) |
| *Infection date and SARS-CoV-2 strain* | 11/08/2021 Presumed Delta (B.1.617.2) | 23/08/2021 Presumed Delta (B.1.617.2) | 18/12/2021 Presumed Omicron (BA.1) |
| *PBMC sample date* | 26/08/2021 | 18/11/2021 | 13/01/2022 |
| *Days post 2nd vaccine* | 163 | 173 | 170 |
| *Days post infection* | 15 | 87 | 26 |

**Supplementary Table 2: Clonally related mAbs isolated from VAIN1, VAIN2 and VAIN3.**

| Donor | Name | Heavy V gene | Heavy CDR3 AA | Heavy CDR3 length | Light V gene | Light CDR3 AA | Light CDR3 length |
|---|---|---|---|---|---|---|---|
| *VAIN1* | V1D_10 | IGHV1-3 | CARGPEMAIVDYFDYW | 16 | IGKV1-5 | CQQYNGYPWTF | 11 |
| | V1D_22 | IGHV1-3 | CARSGGGFLVDYMDVW | 16 | IGKV1-5 | CQQYHGYPWTF | 11 |
| *VAIN2* | V2WT_30 | IGHV3-30 | CARDGKTINMVRGVISGAFDIW | 22 | IGKV1-33 | CQQYDNLPFTF | 11 |
| | V2D_42 | IGHV3-30 | CARDGRTINMVRGVISGAFDIW | 22 | IGKV1-33 | CLQYDILPFTF | 11 |
| | V2WT_11 | IGHV3-30 | CARDGRTITMVRGVISGAFDIW | 22 | IGKV1-33 | CQQYDNLPFTF | 11 |
| | V2WT_23 | IGHV3-30 | CARDGRTITMVRGVISGAFDIW | 22 | IGKV1-33 | CQQYDNLPSTF | 11 |
| | V2D_48 | IGHV3-30 | CARDGTMAPLVPGIMSPAFDIW | 22 | IGKV1-33 | CQQYDNLPPTF | 11 |
| | V2WT_33 | IGHV3-53 | CARDLELAGALDVW | 14 | IGKV1-9 | CQQINSNPPVTF | 12 |
| | V2WT_25 | IGHV3-53 | CARDLELAGGLDIW | 14 | IGKV1-9 | CQQLNSYPPVTF | 12 |
| | V2WT_2 | IGHV4-59 | CARDLAYGEYEGWFDPW | 17 | IGKV1-12 | CQQAYSFPYTF | 11 |
| | V2D_34 | IGHV4-59 | CARDLTYGEYEGWFDPW | 17 | IGKV1-12 | CQQAHSFPYTF | 11 |
| *VAIN3* | V3O_21 | IGHV1-69 | CAIVFGDQSEFDSW | 14 | IGKV3-11 | CQFRSNWPPYTF | 12 |
| | V3O_28 | IGHV1-69 | CAIVFGDQSEFDSW | 14 | IGKV3-11 | CQFRSNWPPYTF | 12 |
| | V3WT_11 | IGHV3-15 | CTTDIYILGVMIEHDAFDSW | 20 | IGKV1-39 | CQQTYTTPAPSF | 12 |
| | V3O_27 | IGHV3-15 | CTTDLYILGVVIEHDAFDIW | 20 | IGKV1-39 | CQQTYFAPALTF | 12 |
| | V3O_37 | IGHV3-53 | CARDFGEMYFDYW | 13 | IGKV3-20 | CQQYGNSPRTF | 11 |
| | V3WT_18 | IGHV3-53 | CARDYGEMYFDFW | 13 | IGKV3-20 | CQQYGGSPRTF | 11 |
| | V3O_26 | IGHV3-66 | CARGFGDQYFDLW | 13 | IGKV1-39 | CQQSYSYPLTF | 11 |
| | V3WT_5 | IGHV3-66 | CARGIGDQYFDLW | 13 | IGKV1-39 | CQQSYSSPLTF | 11 |
| | V3WT_9 | IGHV3-7 | CARGGGHPWYYSGSGSYPPLPKADLDYW | 28 | IGLV1-44 | CVAWDDSLKGSWVF | 14 |
| | V3WT_6 | IGHV3-7 | CARGGGHPWYYSSSGNFPPLPKADLDYW | 28 | IGLV1-44 | CVAWDDSLKGSWVF | 14 |
| | V3O_23 | IGHV4-34 | CARACSGGNCYPRPFDYW | 18 | IGKV1-17 | CLQHNSYPWTF | 11 |
| | V3WT_1 | IGHV4-34 | CARGCSGGICYPKPFDFW | 18 | IGKV1-17 | CLQHNSLPWTF | 11 |

**Supplementary Table 3: Neutralization properties of large-scale expressed nAbs.** mAb are listed based on competition group**.** $IC_{50}$ values are reported in µg/mL. X indicates neutralization not tested.

| Name | IC50 WT | IC50 Beta | IC50 Delta | IC50 Omicron | IC50 BA2 | IC50 BA4 | IC50 D614G | IC50 BA.4 | IC50 XBB | IC50 BQ.1.1 | IC50 BA.2.75.2 | IC50 XBB.1.5 | ACE2 Comp | Specificity | Comp Group | Heavy chain V Gene | Heavy chain CDR3 AA | Light chain V gene | Light chain CDR3 AA |
|---|---|---|---|---|---|---|---|---|---|---|---|---|---|---|---|---|---|---|---|
| VAIN1WT_49 | 0.977 | 2.243 | 2.022 | 14.508 | 11.144 | 6.592 | 6.317 | 10.786 | >50 | >50 | 2.016 | >50 | 24 | RBD | 1 | IGHV4-39 | CARHLEEILPKGVNWFDPW | IGKV1-39 | CQQSYATLPYTF |
| VAIN1D_08 | 0.003 | >50 | 0.001 | >50 | >50 | >50 | 0.001 | >50 | >50 | >50 | >50 | >50 | 0 | RBD | 1 | IGHV1-69 | CARGPPLTRGVARAGQAFDYW | IGKV4-1 | CQQYYSSSITF |
| VAIN1WT_51 | 0.980 | 0.940 | 0.654 | 10.558 | 10.620 | 5.946 | 3.323 | 4.205 | 10.438 | 17.792 | 15.575 | X | 0 | RBD | 1 | IGHV3-30 | CATDSSDPWQNYFDYW | IGKV1-39 | CQQSYSTFEYTF |
| VAIN1WT_25 | 0.130 | 0.044 | 0.080 | >50 | >50 | >50 | X | X | X | X | X | X | 99 | RBD | 1 | IGHV5-10-1 | CARGPSYYHTGRMGDVW | IGKV1-33 | CLQYDSLLGTF |
| VAIN1WT_26 | 0.084 | 0.020 | 0.028 | >50 | >50 | >50 | X | X | X | X | X | X | 98 | RBD | 1 | IGHV1-69 | CACGGYYDTSGYYALDFDSW | IGKV1-5 | CQQFNSYSRTF |
| VAIN2D_06 | 9.274 | 1.457 | 2.700 | >50 | >50 | >50 | X | X | X | X | X | X | 92 | RBD | 1 | IGHV3-23 | CAEPRTGWYVGFDYW | IGLV6-57 | CQSYDSSNHYVF |
| VAIN3WT_20 | 0.082 | 0.147 | 0.079 | 13.395 | 11.448 | 7.948 | X | X | X | X | X | X | 94 | RBD | 1 | IGHV3-11 | CARQKWLRGDFDSW | IGLV6-57 | CQSYDSRNWMF |
| VAIN2D_27 | 0.093 | 0.262 | 0.365 | 1.002 | 1.700 | 0.891 | 0.235 | 1.251 | 0.495 | 0.436 | 0.401 | 3.159 | 78 | RBD | 2 | IGHV3-30 | CARDWGRLVTWFDNW | IGKV1-39 | CQQSYSTPPWTF |
| VAIN3O_12 | 0.135 | 0.044 | 0.053 | 0.529 | 5.745 | 2.475 | 0.008 | 1.849 | 2.176 | 3.916 | 2.337 | 3.076 | 86 | RBD | 2 | IGHV4-61 | CARDLWYYDRSGHYDSDAFDVW | IGLV1-40 | CQSYDSSLTALF |
| VAIN1WT_15 | 0.006 | 0.0005 | 0.014 | 0.001 | 0.002 | 0.004 | 0.023 | 0.001 | >50 | 9.567 | >50 | >50 | 100 | RBD | 2 | IGHV3-53 | CARDLVVYGMDVW | IGKV1-9 | CQQLNSDAISF |
| VAIN2D_12 | 0.010 | 0.002 | 0.014 | 0.001 | 0.003 | 0.001 | 0.008 | 0.003 | 0.004 | 0.004 | 0.005 | 0.002 | 99 | RBD | 3 | IGHV3-66 | CARAYGDRYYFDYW | IGKV1-5 | CQHYGAF |
| VAIN2D_17 | 0.041 | 0.009 | 0.027 | 0.001 | 0.010 | 0.003 | 0.061 | 0.008 | 0.015 | 0.011 | 0.015 | 0.003 | 99 | RBD | 3 | IGHV3-53 | CARDFYRQSGDYPHDFW | IGKV1-33 | CHQYDNLPRTF |
| VAIN3O_37 | 0.002 | 0.0002 | 0.0004 | 0.0002 | 0.0005 | 0.001 | 0.004 | 0.001 | 1.347 | 0.051 | 0.474 | 0.147 | 97 | RBD | 3 | IGHV3-53 | CARDFGEMYFDYW | IGKV3-20 | CQQYGNSPRTF |
| VAIN1WT_14 | 0.014 | 0.005 | 0.027 | >50 | >50 | 0.028 | 0.005 | 0.028 | 0.250 | 0.281 | 0.479 | 0.106 | 97 | RBD | 3 | IGHV3-53 | CARLYNHRRGMDVW | IGKV1-33 | CQQYDNLPITF |
| VAIN2WT_33 | 0.033 | 0.0001 | 0.002 | 0.0002 | 0.0004 | 0.0004 | 0.005 | 0.001 | >50 | 0.103 | 1.699 | 1.211 | 94 | RBD | 3 | IGHV3-53 | CARDLELAGALDVW | IGKV1-9 | CQQINSNPPVTF |
| VAIN2WT_25 | 0.028 | 0.0001 | 0.003 | 0.0002 | 0.0003 | 0.0004 | 0.013 | 0.002 | >50 | 0.219 | 13.184 | >50 | 93 | RBD | 3 | IGHV3-53 | CARDLELAGGLDIW | IGKV1-9 | CQQLNSYPPVTF |
| VAIN3WT_05 | 0.006 | 0.001 | 0.003 | 0.001 | 0.002 | 0.001 | 0.002 | 0.001 | >50 | 2.527 | 2.477 | >50 | 99 | RBD | 3 | IGHV3-66 | CARGIGDQYFDLW | IGKV1-39 | CQQSYSSPLTF |
| VAIN1D_30 | 0.005 | 0.001 | 0.007 | 0.0001 | 0.001 | 0.438 | 0.006 | 0.271 | >50 | 1.865 | >50 | >50 | 98 | RBD | 3 | IGHV3-64 | CVKGGIQLWPTGDSW | IGKV1-5 | CQQYKSYPWTF |
| VAIN3O_23 | 0.005 | 0.001 | 0.0003 | 0.001 | 0.001 | >50 | 0.001 | >50 | >50 | >50 | >50 | >50 | 97 | RBD | 3 | IGHV4-34 | CARACSGGNCYPRPFDYW | IGKV1-17 | CLQHNSYPWTF |
| VAIN2WT_23 | 0.004 | >50 | 0.0005 | >50 | >50 | >50 | 0.004 | >50 | >50 | >50 | >50 | >50 | 89 | RBD | 3 | IGHV3-30 | CARDGRTITMVRGVISGAFDIW | IGKV1-33 | CQQYDNLPSTF |
| VAIN2D_48 | 0.034 | >50 | 0.002 | >50 | >50 | >50 | 0.034 | >50 | >50 | >50 | >50 | >50 | 90 | RBD | 3 | IGHV3-30 | CARDGTMAPLVPGIMSPAFDIW | IGKV1-33 | CQQYDNLPPTF |
| VAIN1D_25 | 0.023 | 0.002 | 0.013 | 0.001 | 0.003 | 1.191 | 0.005 | 0.756 | >50 | >50 | >50 | X | 99 | RBD | 3 | IGHV3-53 | CARDLAPVGLMDVW | IGKV1-27 | CQNYNSDPPWTF |
| VAIN1WT_43 | 0.121 | 0.594 | 0.660 | 0.117 | 3.754 | 1.287 | 2.210 | 6.019 | 10.646 | 34.975 | 13.252 | X | 0 | RBD | 3 | IGHV3-33 | CARDEGAVVTHMDYW | IGKV1-39 | CQQSYNTPPWTF |
| VAIN1WT_39 | 0.001 | 2.460 | 0.002 | >50 | >50 | >50 | X | X | X | X | X | X | 99 | RBD | 3 | IGHV5-10-1 | CARQSGDYFDYLLAYFDLW | IGKV3-20 | CQQYSSSPPGYTF |
| VAIN2WT_11 | 0.005 | >50 | 0.003 | >50 | >50 | >50 | X | X | X | X | X | X | 81 | RBD | 3 | IGHV3-30 | CARDGRTITMVRGVISGAFDIW | IGKV1-33 | CQQYDNLPFTF |
| VAIN2WT_18 | 10.565 | 21.841 | 8.213 | >50 | >50 | >50 | X | X | X | X | X | X | 18 | RBD | 3 | IGHV3-53 | CARVLPYGDNYFDFW | IGKV3-11 | CQQLFTF |
| VAIN2D_02 | 0.004 | 0.130 | 0.005 | >50 | 39.604 | 1.000 | X | X | X | X | X | X | 99 | RBD | 3 | IGHV3-66 | CARNVWDDAFDLW | IGKV1-9 | CQLLNSYPPGTF |
| VAIN1WT_41 | 0.051 | 0.044 | 0.089 | 0.120 | 1.498 | 1.431 | X | X | X | X | X | X | 95 | RBD | 3 | IGHV4-39 | CARTAPYYYDRSGYYQKEEYFQRW | IGKV1-5 | CQQYNNYPRTF |
| VAIN2WT_03 | 0.006 | >50 | 0.010 | >50 | >50 | >50 | X | X | X | X | X | X | 0 | RBD | 3 | IGHV3-30-3 | CARDGTMAPLVPGIMSPAFDIW | IGKV1-33 | CQQYDNLPPTF |
| VAIN3WT_01 | 0.004 | 0.001 | 0.0003 | 0.001 | 0.002 | >50 | 0.002 | >50 | >50 | >50 | >50 | >50 | 0 | RBD | 3.5 | IGHV4-34 | CARGCSGGICYPKPFDFW | IGKV1-17 | CLQHNSLPWTF |
| VAIN2WT_01 | 0.002 | 2.352 | 0.005 | 0.482 | 0.101 | >50 | 0.001 | >50 | >50 | 31.732 | 6.108 | X | 99 | RBD | 3.5 | IGHV3-53 | CARESYAVATIGKEYRMDVW | IGKV3-11 | CQQRSNWPWGF |
| VAIN1WT_09 | 0.012 | >50 | 0.001 | >50 | >50 | >50 | X | X | X | X | X | X | 76 | RBD | 3.5 | IGHV1-8 | CARGGRYCDITSCYSRGWLDPW | IGLV1-51 | CGTWDAGLSVVF |
| VAIN1D_03 | 0.017 | >50 | 0.002 | 1.587 | >50 | >50 | X | X | X | X | X | X | 0 | RBD | 3.5 | IGHV1-2 | CARDQSFSMVRGDTDHW | IGKV1-39 | CQQSYRTPALSF |
| VAIN1WT_02 | 0.088 | 0.236 | 0.230 | 0.923 | 2.235 | 3.429 | X | X | X | X | X | X | 57 | RBD | 3.5 | IGHV1-46 | CARAGVAPDHSHPFDPW | IGKV1-13 | CQQFNTYLSITF |
| VAIN1WT_06 | 0.225 | 0.276 | 0.306 | 0.310 | 1.326 | 0.620 | 2.334 | 4.325 | 1.289 | 4.834 | 7.452 | 1.551 | 72 | RBD | 4 | IGHV1-18 | CARGPWGYCSGTSCYYWTPDNPAYSYHGMDVW | IGKV2-24 | CMQATQFPHTF |
| VAIN1WT_18 | 0.070 | 0.094 | 0.063 | 0.198 | 0.447 | 0.114 | 0.180 | 0.613 | 0.288 | 0.538 | 0.539 | 0.678 | 0 | RBD | 4 | IGHV5-51 | CARQFCGGGCHFDYW | IGKV1-5 | CQQYNSYSRTF |
| VAIN1WT_32 | 0.133 | 0.090 | 0.103 | 0.089 | 0.233 | 0.212 | 0.265 | 0.261 | 0.713 | 2.042 | 2.976 | 0.843 | 28 | RBD | 4 | IGHV5-51 | CARSDTSNYFDYW | IGKV1-5 | CQQYNGYSF |
| VAIN1WT_50 | 0.251 | 0.160 | 0.245 | 0.210 | 1.036 | 0.494 | 2.340 | 1.597 | 2.329 | 3.862 | 1.907 | 2.044 | 0 | RBD | 4 | IGHV1-69 | CAKGGGYSYGHHYNWFFDVW | IGKV4-1 | CQQYFSTPPWTF |
| VAIN1D_19 | 0.139 | 0.034 | 0.075 | 0.065 | 0.268 | 0.160 | 0.182 | 0.067 | 0.142 | 1.392 | 1.093 | 0.128 | 4 | RBD | 4 | IGHV5-51 | CARTLQTNWLDHW | IGKV1-5 | CQQFNTYWTF |
| VAIN2WT_29 | 0.102 | 0.040 | 0.320 | 0.332 | 2.220 | 0.557 | 0.102 | 0.557 | 4.580 | 4.338 | 1.787 | 1.244 | 18 | RBD | 4 | IGHV5-51 | CARRTSSAGLPCLDVW | IGKV1-39 | CQQSFSTLALTF |
| VAIN2D_28 | 0.083 | 3.039 | 0.540 | 3.786 | 2.133 | 1.300 | 5.469 | 2.688 | 0.744 | 1.007 | 2.138 | 2.637 | 60 | RBD | 4 | IGHV3-48 | CARDTGFWSGHYPAQFDYW | IGKV1-33 | CQQYNTPPPVTF |
| VAIN2D_25 | 0.078 | 0.011 | 0.029 | 0.006 | 0.010 | 0.023 | 0.550 | 0.153 | 0.278 | >50 | 0.183 | 1.016 | 54 | RBD | 4 | IGHV5-51 | CARRDSGSYSGYFDLW | IGLV1-44 | CAAWDDSLNGVVF |
| VAIN3O_09 | 0.016 | 0.001 | 0.002 | 0.001 | 0.001 | 0.006 | 0.006 | 0.004 | 0.038 | >50 | 0.034 | 0.028 | 97 | RBD | 4 | IGHV4-39 | CTRMAVQWGWYGAIDYW | IGLV3-21 | CQVWESIYDLPWVF |
| VAIN3O_15 | 0.015 | 0.007 | 0.0002 | 0.006 | 0.007 | 0.002 | 0.013 | 0.001 | 0.041 | >50 | 0.009 | 0.049 | 0 | RBD | 4 | IGHV4-31 | CARGIPDSAVNSW | IGLV1-40 | CQSYDSSMSGPVF |
| VAIN1D_05 | 0.181 | 0.077 | 0.042 | 0.016 | 0.037 | 0.064 | 0.144 | 0.229 | 0.766 | >50 | 0.233 | 0.646 | 0 | RBD | 4 | IGHV1-69 | CARDMIEAPLYGMDVW | IGKV1-39 | CQQSYRTPALSF |
| VAIN1D_22 | 0.096 | 0.007 | 0.008 | 0.004 | 0.007 | 0.004 | 0.017 | 0.003 | >50 | >50 | 1.785 | >50 | 0 | RBD | 4 | IGHV1-3 | CARSGGGFLVDYMDVW | IGKV1-5 | CQQYHGYPWTF |
| VAIN3WT_07 | 0.019 | 0.002 | 0.025 | 0.001 | 0.008 | 0.005 | 0.001 | 0.001 | >50 | >50 | 0.093 | >50 | 96 | RBD | 4 | IGHV3-15 | CVTDLFVQGVILEHDAFDIW | IGKV1-39 | CQQCYITPAPTF |
| VAIN3O_03 | 0.164 | 0.008 | 0.075 | 0.020 | 0.007 | 0.065 | 0.033 | 0.030 | >50 | >50 | 0.240 | >50 | 0 | RBD | 4 | IGHV3-9 | CVKEKIRLREGEGMDVW | IGLV3-21 | CQVWESSSDHVVF |
| VAIN3WT_13 | 0.007 | 0.010 | 0.024 | 0.857 | 0.834 | 0.688 | 0.004 | 0.272 | >50 | >50 | >50 | >50 | 95 | RBD | 4 | IGHV1-18 | CARDSSVAGSLDYW | IGLV2-14 | CSSYTRTIYIF |
| VAIN1D_10 | 0.002 | 0.001 | 0.003 | 0.008 | 0.002 | 0.002 | 0.001 | 0.000 | >50 | >50 | >50 | >50 | 9 | RBD | 4 | IGHV1-3 | CARGPEMAIVDYFDYW | IGKV1-5 | CQQYNGYPWTF |
| VAIN3WT_02 | 0.619 | 0.018 | 12.037 | 0.019 | 0.156 | >50 | 0.085 | >50 | >50 | >50 | 0.002 | >50 | 0 | RBD | 4 | IGHV1-46 | CARSHSVTLAGVDYW | IGLV7-43 | CLLYYEGAWVF |
| VAIN2D_43 | 48.000 | 31.281 | 48.000 | >50 | >50 | >50 | 48.000 | >50 | >50 | >50 | >50 | >50 | 19 | RBD | 4 | IGHV3-30 | CAKDRSWFGEFQDDNYYYAMDVW | IGKV1-33 | CQQYDNLPLSF |
| VAIN3WT_21 | 0.019 | 0.022 | 0.454 | 0.019 | 0.019 | 6.590 | X | X | X | X | X | X | 78 | RBD | 4 | IGHV3-30 | CVSFSASGTYPLYFDYW | IGKV1-33 | CHQYDNLPQTF |
| VAIN3O_08 | 0.119 | 0.016 | 0.730 | 0.044 | 0.098 | 8.565 | X | X | X | X | X | X | 10 | RBD | 4 | IGHV3-9 | CAKDLHSGSYLMAHDAFDVW | IGLV1-51 | CGTWDSSLSAGVS |
| VAIN1D_02 | 3.956 | 3.792 | 2.104 | 1.573 | 3.262 | 4.502 | X | X | X | X | X | X | 23 | RBD | 4 | IGHV3-23 | CAKEGPYYCSSSSCYVGIFDFW | IGKV1-13 | CQQFQRYSFTF |
| VAIN1D_21 | 0.308 | 0.017 | 0.202 | 0.125 | 0.260 | 0.406 | X | X | X | X | X | X | 0 | RBD | 4 | IGHV5-51 | CARSRSSDYFDYW | IGKV1-5 | CQQYKSYSKTF |
| VAIN2WT_12 | 8.648 | 11.377 | 5.392 | 13.725 | 13.283 | 7.194 | X | X | X | X | X | X | 26 | RBD | 4 | IGHV3-30 | CAKGAGPYCSAGDCLRQGLDYW | IGKV1-33 | CQQYDNLPLTF |
| VAIN1WT_36 | 0.001 | >50 | 0.005 | 0.339 | >50 | >50 | X | X | X | X | X | X | 83 | RBD | 4 | IGHV3-21 | CARNLGASAGVNIIHYFYHYMDVW | IGLV2-23 | CCSYAGTGSPWVF |
| VAIN2D_36 | 15.179 | 49.955 | >50 | 9.002 | 9.002 | 11.919 | X | X | X | X | X | X | 60 | NTD | 5 | IGHV4-34 | CARGLLRIFIPVGGVGGFDSW | IGKV3-15 | CQQYKYWPPITF |
| VAIN2D_16 | 0.110 | >50 | >50 | 24.364 | >50 | 10.860 | X | X | X | X | X | X | 54 | NTD | 5 | IGHV4-34 | CARYLQWLVPPGFDPW | IGKV1-39 | CQQSYSTPRTF |
| VAIN1D_06 | 0.331 | 0.013 | 0.018 | >15 | 0.0003 | 0.002 | 0.023 | 0.000 | >50 | 0.004 | >50 | >50 | 52 | NTD | 6 | IGHV1-24 | CATSPAVAGTMEDYSYYYGMDVW | IGKV2-24 | CMQATQFPITF |
| VAIN3WT_06 | 3.043 | 0.157 | >50 | 0.037 | 20.162 | 37.554 | 1.000 | 27.488 | 31.302 | 46.588 | >50 | >50 | 0 | NTD | 6 | IGHV3-7 | CARGGGHPWYYSSSGNFPPLPKADLDYW | IGLV1-44 | CVAWDDSLKGSWVF |
| VAIN3O_10 | 0.772 | 0.004 | >50 | 0.004 | >50 | >50 | X | X | X | X | X | X | 49 | NTD | 6 | IGHV3-7 | CTRDASNWGEWLRNSYYYYMDVW | IGLV1-44 | CAAWDEGLNTWMF |
| VAIN1WT_13 | 0.033 | 1.138 | 0.045 | 2.185 | 13.421 | 13.950 | X | X | X | X | X | X | 0 | NTD | x | IGHV3-7 | CARDAYESSGWPVAFDIW | IGKV3-20 | CQQYGSHF |
| VAIN1WT_46 | 0.014 | >50 | 14.084 | >50 | >50 | >50 | X | X | X | X | X | X | 20 | NTD | x | IGHV4-39 | CTRYLRGYSEDFPFDIW | IGKV1-17 | CLQYNSYPHTF |

**Supplementary Table S4: Mutations present in variant Spikes.** Unique mutations are highlighted in red.

| Variant | Spike Mutations |
|---|---|
| **Beta (B.1.351)** | L18F, D80A, D215G, (Del 241-243), K417N, E484K, N501Y, D614G, A701V |
| **Delta (B.1.617.2)** | T19R, G142D, 156del, 157del, R158G, L452R, T478K, D614G, P681R, D950N |
| **Omicron (BA.1)** | A67V, 69-70del, T95I, GVYY142-145D, NL211-212I, ins214EPE, G339D, S371L, S373P, S375F, K417N, N440K, G446S, S477N, T478K, E484A, Q493R, G496S, Q498R, N501Y, Y505H, T547K, D614G, H655Y, N679K, P681H, N764K, D796Y, N856K, Q954H, N969K, L981F |
| **Omicron (BA.2)** | T19I, LPPA24-27S, G142D, V213G, G339D, S371F, S373P, S375F, T376A, D405N, R408S, K417N, N440K, S477N, T478K, E484A, Q493R, Q498R, N501Y, Y505H, D614G, H655Y, N679K, P681H, N764K, D796Y, Q954H, N969K |
| **Omicron (BA.4)** | T19I, LPPA24-27S, Del 69-70, G142D, V213G, G339D, S371F, S373P, S375F, T376A, D405N, R408S, K417N, N440K, L452R, S477N, T478K, E484A, F486V, Q498R, N501Y, Y505H, D614G, H655Y, N679K, P681H, N764K, D796Y, Q954H, N969K |
| **Omicron (BA.2.75.2)** | T19I, LPPA24-27S, G142D, K147E, W152R, F157L, I210V, V213G, G257S, G339H, S371F, S373P, S375F, T376A, D405N, R408S, K417N, N440K, G446N, N460K, S477N, T478K, E484A, Q498R, N501Y, Y505H, D614G, H655Y, N679K, P681H, N764K, D796Y, Q954H, N969K |
| **Omicron (BQ1.1)** | T19I, LPPA24-27S, H69del, V70del, V213G, G142D, G339D, R346T, S371F, S373P, S375F, T376A, D405N, R408S, K417N, N440K, K444T, L452R, N460K, S477N, T478K, E484A, F486V, Q498R, N501Y, Y505H, D614G, H655Y, N679K, P681H, N764K, D796Y, Q954H, N969K |
| **Omicron (XBB)** | T19I, LPPA24-27S, V83A, G142D, Del144, H146Q, Q183E, V213E, G252V, G339H, R346T, L368I, S371F, S373P, S375F, T376A, D405N, R408S, K417N, N440K, V445P, G446S, N460K, S477N, T478K, E484A, F486S (**XBB1.5:** F486P**)**, F490S, Q498R, N501Y, Y505H, D614G, H655Y, N679K, P681H, N764K, D796Y, Q954H, N969K |
